# The diversification of *Pterocarpus* (Leguminosae: Papilionoideae) was influenced by biome-switching and infrequent long-distance dispersal

**DOI:** 10.1101/2021.05.14.444160

**Authors:** Rowan J. Schley, Ming Qin, Mohammad Vatanparast, Panagiota Malakasi, Manuel de la Estrella, Gwilym Lewis, Bente Klitgård

## Abstract

**Aim:** Phenotypes which evolved for dispersal over ecological timescales may lead to significant macroevolutionary consequences, such as infrequent long-distance dispersal and diversification in novel biomes. We aimed to reconstruct the phylogenetic history of *Pterocarpus* (Leguminosae/ Fabaceae) to assess whether seed dispersal phenotypes and biome switching explain the current biogeographical patterns of this group.

**Location:** Pantropical

**Taxon:** The Pterocarpus clade, particularly *Pterocarpus* (Leguminosae/Fabaceae)

**Methods:** We sequenced ~300 nuclear loci captured using *Angiosperms*-353, a genomic ‘bait set’ for flowering plants, from which we generated a time-calibrated phylogenomic tree. To corroborate this, we also generated a time-calibrated phylogenetic tree from data-mined Sanger-sequencing data. We then collated distribution data and fruit dispersal morphology traits to compare trait-dependent and trait-independent biogeographical models, allowing us to assess whether dispersal traits influenced the spatio-temporal evolution of *Pterocarpus*. Finally, using the results of these model tests, we estimated the ancestral ranges and biomes of *Pterocarpus* species to better understand their biogeographical history, and assessed the degree and direction of biome switching over the course of *Pterocarpus*’ diversification history.

**Results:** We recovered well-supported phylogenetic relationships within *Pterocarpus*, within which there were two subclades – one Neotropical and the other Palaeotropical. Our divergence date estimates suggested that *Pterocarpus* largely diversified from around 12 Ma, during the Miocene.

Trait-dependent biogeographical models were rejected for both range and biome evolution within *Pterocarpus*, but models parameterising dispersal were supported. *Pterocarpus*’ ancestral node shared a range across the new-world and old-world tropics, followed by divergence into two clades, one palaeotropical and one neotropical. Biome switching occurred most frequently into rainforest and grassland.

**Main conclusions:** Overall, our analyses suggest that *Pterocarpus* underwent infrequent cross-continental dispersal and establishment into novel biomes. While this was minimally impacted by fruit dispersal syndromes, biome switching precipitated by long-distance dispersal and environmental change have played an important role in diversification within *Pterocarpus* since the Miocene.

## Introduction

Dispersal is a major process structuring the geographic ranges of organisms, both at ecological and evolutionary timescales (Brown et al., 1996, Sanmartín & Ronquist, 2004). Indeed, adaptations which evolved to promote dispersal locally may end up being significant evolutionary innovations, influencing the biogeographical fate of lineages by allowing dispersal across continents and subsequent diversification over millions of years (Klaus & Matzke, 2020).

While vicariance has a major role in explaining biogeographical patterns, the role of dispersal was seen as being somewhat less significant (Sanmartín, 2012). However, there is now ample evidence that dispersal, and especially trans-oceanic dispersal, plays a major role in the biogeographical dynamics of a great number of terrestrial lineages (De Queiroz, 2005, Gillespie et al., 2012, Harris et al., 2018). This is especially evident in plants due to the wide range of traits that they have evolved to promote dispersal of their fruits, seeds and spores (Cousens et al., 2008). Seeds in particular have developed a wide range of morphological traits which allow dispersal over long distances, including wings and other aerodynamic structures (Cain et al., 2000), long dormancy periods (van der Pijl, 1982), thick seed coats and air pockets which render the seeds buoyant, thus facilitating dispersal across oceans (Bellot et al., 2020, Renner, 2004). Multiple recent studies have highlighted that the biogeographical distributions of modern plant lineages are largely explained by traits which they evolved for effective dispersal in their ecological context (e.g. Podocarpaceae (Klaus & Matzke, 2020); Annonaceae (Onstein et al., 2019)) using trait-dependent biogeographical modes. Indeed, it has been well established that a wide range of phenotypic traits have a significant impact on macroevolutionary dynamics (e.g. Rabosky et al., 2014).

Given their superlative biological diversity, the tropics are one of the most important places to examine the impact of dispersal on biogeography and diversification. Dispersal is particularly important for terrestrial organisms in the tropics because they are subject to both geographical and ecological constraints to their distribution. The tropics are largely separated into three zones (Neotropical, African and Asian/Australasian) by large areas of ocean, bounded to the north and south by increasingly unsuitable climates with increasing distance from the equator (Corlett & Primack, 2011, Morley, 2000). There is ample evidence of disjunct ranges across continents within tropical organisms, many of which likely result from dispersal, particularly in plants (e.g., Arecaceae (Eiserhardt et al., 2011), Podocarpaceae (Klaus & Matzke, 2020), Leguminosae/Fabaceae (Schrire et al., 2005, Vatanparast et al., 2013), Vitaceae (Nie et al., 2012)).

Following dispersal, plants may subsequently encounter powerful selective pressures in contrasting habitats to those in which they evolved. This can drive adaptation, and eventually speciation, to these novel environments (Waser & Campbell, 2004). Biomes which span climatic extremes are found in relatively close proximity within the tropics, being defined by their ecological community composition (Pennington & Dick, 2004, Sanmartín, 2014) as well as by the physiognomy of their dominant plant species (Pennington & Ratter, 2006, Woodward et al., 2004). These biomes are often seen as independent evolutionary arenas (Hughes et al., 2013, Nürk et al., 2020, e.g. Pennington et al., 2009, Ringelberg et al., 2020). This biome concept reflects that environmental pressures select for species with similar functional attributes (Echeverría-Londoño et al., 2018), further implying that colonisation of contrasting biomes can promote speciation (Pennington & Dick, 2010). For example, many genera with species typical of seasonally dry tropical forest and savanna also contain species from rainforest (Pennington & Ratter, 2006, Pennington et al., 2000), suggesting that biome switching has driven ecological speciation.

The relative invasibility of certain biomes by immigrants may be higher than in others. Globally, lowland tropical ecosystems such as rainforest and savanna tend to be most permeable to colonisation by taxa from other biomes (Dexter et al., 2015), acting as ‘lineage sinks’ (Donoghue & Edwards, 2014, Pennington & Hughes, 2014) because plant communities in these environments experience high species turnover through exposure to drought and fire (da Costa et al., 2010, Pennington & Lavin, 2016). Such an imbalance in the direction of biome switching in tropical plant taxa is also likely to be an outcome of the relative size of the adaptive peak to be overcome by immigrants, coupled with the pre-existence of adaptive traits (Donoghue & Edwards, 2014). Again, drought and fire are likely to be modulators of the shape of such adaptive peaks, requiring specialised phenotypes to survive them (De Micco & Aronne, 2012, Midgley & Bond, 2013) and constraining plant evolution in the tropics (Olmstead, 2013).

Pantropical clades which occur in different biomes are an ideal system to study the biogeographical processes outlined above. The legume family (Leguminosae/Fabaceae) is replete with such clades, present in every terrestrial biome in the tropics (Lewis, G. P. et al., 2005) and dominant in many of them (de la Estrella et al., 2017, Gentry, 1988, ter Steege et al., 2013, ter Steege et al., 2020). The genus *Pterocarpus* Jacq. belongs to the largest subfamily of legumes (Papilionoideae) and consists of around 33 species (Klitgård & Lavin, 2005, Lewis, G. P., 1987, Rojo, 1972), many of which are important timber species or are used in traditional medicine (Saslis-Lagoudakis et al., 2011). This genus is diagnosed by either winged or corky (coriaceous) fruits adapted for dispersal by wind or water, respectively, and this is likely one reason why *Pterocarpus* is found pantropically (Klitgård & Lavin, 2005). *Pterocarpus* forms a part of the broader, pantropical ‘Pterocarpus clade’ which contains 22 genera and *~*200 species, of which most are trees (Klitgård et al., 2013).

Given its pantropical distribution, its presence in many contrasting biomes and its diversity of dispersal syndromes, *Pterocarpus* is an ideal system for evolutionary studies of biogeography, biome-switching and dispersal within the tropics (Klitgård et al., 2013). Accordingly, we aim to examine the impact of dispersal traits on the diversification of *Pterocarpus* to provide insights into how trait evolution can structure the biogeographical distribution of species, and so influence the evolution of tropical diversity. We also aim to infer the patterns of biome switching which may result from such dispersal events, to help clarify how shifts between biomes may accumulate or generate diversity. To do this, we will reconstruct the relationships between species within the Pterocarpus clade using novel next-generation sequencing techniques, and infer a time-calibrated phylogeny for *Pterocarpus* using multiple approaches. Following this, we will test whether trait-dependent or trait-independent biogeographical models best explain extant biogeographical patterns, both in terms of continental realm and biome, and assess whether transitions between certain biomes are more frequent than others. As such, we predict that:

1. Extant biogeographical patterns in *Pterocarpus* were significantly influenced by their seed dispersal traits, and hence that a trait-dependent model will best describe their spatiotemporal evolution.
2. Biome shifts will be more frequent into rainforests and savannas, given their ecologically dynamic nature and resulting high invasibility by taxa dispersing into them.

## Materials and Methods

### Taxon sampling

Novel phylogenomic data were generated using target capture sequencing for 26/33 accepted *Pterocarpus* species (75%), which were sampled across the distributional range of the genus. In total, 27 accessions were sampled within *Pterocarpus*, detailed in Supporting Information, Appendix S1, Table S1.1. A further 22 outgroup taxa from the 21 other genera belonging to the Pterocarpus clade within which *Pterocarpus* is nested were sampled for phylogenetic analysis. In addition, Sanger sequence data from one nuclear locus (nrITS) and four plastid loci (*mat*K, *ndhF-rpl*32, *rbcl, trn*L-*trn*F) were downloaded from NCBI GenBank https://www.ncbi.nlm.nih.gov/genbank/ (Appendix S1, Table S1.2) for comparison between phylogenetic dating methods. These data represented 28/33 *Pterocarpus* species, along with 8 outgroup species from the Pterocarpus clade.

A species list of all accepted *Pterocarpus* taxa was compiled using IPNI (IPNI, 2020) and a generic monograph of *Pterocarpus* (Rojo, 1972) to ensure that taxonomically accepted species were included in analyses, and to account for synonyms and infraspecific taxa. Voucher specimens were examined, and their determination updated as appropriate. Newly described and recently reinstated species from within the *P. rohrii* complex were also sampled (Klitgård et al., *in prep)*. Leaf material for DNA extraction was acquired from both silica material *(N=* 1) and herbarium specimens (*N*= 48) collected from the BM, K and MO herbaria.

### Library preparation and sequencing

DNA was extracted from 20 mg of dried leaf material with the CTAB method (Doyle & Doyle, 1987), following which DNA concentrations were measured using a Quantus fluorometer (Promega, Wisconsin, USA). Libraries were prepared using the NEBNext® Ultra™ II DNA Kit (New England Biolabs, Massachusetts, USA). Samples with high molecular weight DNA (>1000 bp, measured with a TapeStation 4200 (Agilent Technologies, California, USA)) were sheared with a Covaris focussed ultrasonicator M220 (Covaris, Massachusetts, USA). Following this, end-preparation and Illumina adaptor ligation were undertaken according to the NEBNext protocol, including a 400bp size-selection step with Agencourt Ampure XP magnetic beads (Beckman Coulter, California, USA). Libraries containing a range of insert sizes were then amplified using PCR, following the protocol outlined in Appendix S1, Table S1.2. Targeted bait capture was performed using the MyBaits protocol (Arbor Biosciences, Michigan, USA) to target 353 phylogenetically informative nuclear genes with *Angiosperms*-353, a bait kit designed for all Angiosperms. The final library pools were sequenced using a paired-end 150bp run on the Illumina HiSeq platform by Macrogen 154 Inc. (Seoul, South Korea).

### Quality filtering, read assembly and alignment

DNA sequencing reads were quality-checked with FASTQC v0.11.3 (Andrews, 2010) and were trimmed using Trimmomatic v.0.3.6 (Bolger et al., 2014) to remove adapter sequences and quality-filter reads. Trimmomatic settings permitted <4 mismatches, a palindrome clip threshold of 30 and a simple clip threshold of 6. Bases with a quality score <28 and reads shorter than 36 bases long were removed from the dataset. Following quality-filtering, loci were assembled using SPADES v3.11.1 (Bankevich et al., 2012) by the HybPiper pipeline v1.2 (Johnson et al., 2016) with a coverage cut-off of 8x. All loci with <50% recovery among taxa were removed, and potentially paralogous loci were removed using the Python (Python Software Foundation, 2010) script *‘paralog_investigator.py’*, distributed with the HybPiper pipeline. Recovery of different loci was visualised using the *‘gene_recovery_heatmap.R’* script distributed with HybPiper.

Following this, targeted loci were aligned by gene region (excluding those with potential paralogs) using 1,000 iterations in MAFFT (Katoh & Standley, 2013) with the ‘—*adjustdirectionaccurately’* option to incorporate reversed sequences. These alignments were then cleaned using the □*automated1* flag in TRIMAL (Capella-Gutiérrez et al., 2009), following which they were visually inspected for poor sequence quality using *Geneious* v. 8.1.9 (https://www.Geneious.com). This was done in order to remove taxa with mostly missing data and to prevent the inclusion of poorly recovered loci into the dataset, resulting in 303 refined alignments.

The nrITS and four plastid Sanger sequencing loci downloaded from NCBI GenBank were refined to include 68 sequences belonging to 36 species from nine genera (GenBank numbers for these sequences are listed in Appendix S1, Table S1.3). These data were aligned using MAFFT with the same parameters as above. These sequences were also subject to visual data inspection and trimming in Geneious to minimise missing sequence data where multiple accessions of a species were present.

### Phylogenomic inference and divergence dating

Three hundred and three gene trees were inferred using RAxML v.8.0.26 (Stamatakis, 2014) with 1,000 rapid bootstrap replicates and the GTRCAT model of nucleotide substitution. A species tree was generated based on the best-scoring RAxML trees using ASTRAL v.5.6.1 under the default parameters, with monophyly not enforced (‘*-a*’ flag) (Zhang, C. et al., 2018). To compare between phylogenetic methods, we then concatenated all 303 target capture locus alignments into a single alignment using AMAS (Borowiec, 2016), and inferred a phylogenetic tree using RAxML HPC2 (Stamatakis, 2014) on the CIPRES web portal ((Miller et al., 2010), https://www.phylo.org/). Inference was performed using 1,000 rapid bootstrap replicates and the GTRCAT model of nucleotide substitution.

MCMCtree, from the PAML package (Yang, 2007), was used to date our phylogenomic tree. This program allows Bayesian divergence time estimation from large phylogenomic datasets based on a pre-determined phylogenetic tree (Dos Reis & Yang, 2013). As such, the ASTRAL tree was rooted with an appropriate outgroup *(Cascaronia astragalina)* and the tree’s branch lengths were removed with the R (R Development Core Team, 2013) package ‘PhyTools’ (Revell, 2012). This rooted, branchless tree and the concatenated 303-locus target capture alignment were then used as inputs for MCMCtree. Fossil calibrations (listed in Table 1) were added using skew-normal priors as uncertainty is distributed asymmetrically across the mean age in fossil calibrations (Ho & Phillips, 2009) using the *R* package ‘MCMCtreeR’ (Puttick, 2019).

**Table 1:**
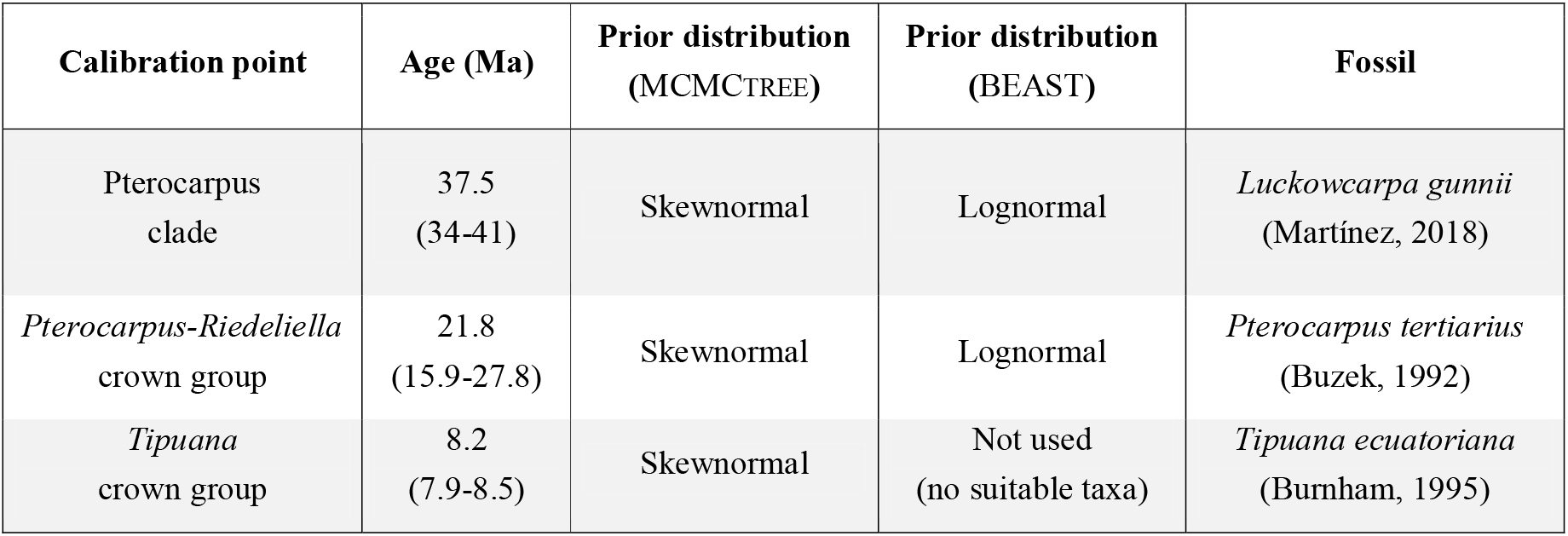
Fossils used to generate a time-calibrated phylogenomic tree of the Pterocarpus clade in MCMCtree and a dated phylogenetic tree in BEAST v.1.8.0. Skewnormal calibration priors were used for fossil ages in MCMCtree to provide a minimum age estimate, and the corresponding Lognormal priors were used for the BEAST analysis. Mean ages are shown for calibrations, with 95% confidence intervals in parentheses. All ages are in millions of years (Ma).

The MCMCtree analysis was run by first calculating the gradient (b) and Hessian (h) parameters using the HKY substitution model, from which the posteriors of divergence times and rates were estimated using two independent runs of 20 million generations to ensure MCMC convergence. A third, identical run with only priors was performed and compared to the runs with data to assess whether prior settings were adequate. MCMC convergence for posterior estimation was checked using Tracer (Rambaut et al., 2015), as well as in the R packages ‘ape’ and ‘bppr’ (Angelis & Dos Reis, 2015, Paradis & Schliep, 2019). The final time-calibrated phylogenetic tree estimated with MCMCtree was then plotted with ‘MCMCtreeR’.

The GenBank dataset of five Sanger-sequenced loci was used to perform corroborative divergence time estimation in BEAST v.1.8.0 (Drummond & Bouckaert, 2015) on the CIPRES web portal. Substitution rate models were chosen for each of the five loci using AlCc in JModelTest (Darriba et al., 2012), and are outlined in Appendix S1, Table S1.4. Tree model choice was based on path sampling analysis output from BEAST, using 10 million generations with 30 path steps. Two fossil calibration points from the MCMCtree analysis were used to provide date estimates in BEAST (Table1). BEAST comprised three independent runs of 50 million generations which were combined using Logcombiner v.1.8.2 with a burn-in level of 10%, using a birth-death tree model and a lognormal clock model. Convergence of runs was assessed using effective sample size (ESS) of all parameters in Tracer v. 1.6 with a threshold value of 200. Final ultrametric trees were generated in TreeAnnotator v1.8.2 and visualised using FIGTree v1.4.4 (Rambaut, 2014). Further information on BEAST analysis is available in Appendix S1, Supplementary Methods.

### Biogeographical analyses

We tested trait-dependent biogeographical models in the R package ‘BioGeoBEARS’ (Matzke, 2013) to understand the relationships between seed dispersal traits, biome switching and the spatio-temporal evolution of *Pterocarpus*. Fruit morphology traits were collected from the first monograph of *Pterocarpus* (Rojo, 1972), a taxonomic revision circumscribing new species within the *P. rohrii* complex (Klitgård *et al. in prep)*, and taxonomic treatments of various *Pterocarpus* species (de Candolle, 1825, Klitgård et al., 2000, Zamora, 2000). Since no fruiting specimen of *P. rohrii* var. *rubiginosus* exists, it was inferred to be winged since it belongs to the *P. rohrii* species complex. Fruit dispersal traits were then plotted on the *Pterocarpus* ASTRAL tree with biome and realm for each species using the *dotplot()* function in the R package ‘PhyTools’.

Geographical distribution data for non-cultivated *Pterocarpus* specimens were collected from GBIF (www.GBIF.org (06 November 2019), GBIF Occurrence Download https://doi.org/10.15468/dl.tr3h2y) based on the curated taxonomic name list described above. This dataset was then further refined using the R package ‘CoordinateCleaner’ ((Zizka et al., 2019); https://github.com/ropensci/CoordinateCleaner) and the input for our *BioGeoBears* analysis was produced from these data with Alex Zizka’s ‘biogeography in R’ scripts (https://github.com/azizka/Using_biodiversity_data_for_biogeography). These records and the specimens underpinning them were then examined in detail by taxonomic experts on the *Pterocarpus* clade (B. B. Klitgaard and G. P. Lewis) to further remove records which were mis-identified.

Based on these data, we assigned species to biogeographical regions using the *wwfLoad()* function in ‘speciesgeocodeR’ (Töpel et al., 2017), based on the WWF ecoregions of Olson et al. (2001). These ecoregions represented five tropical terrestrial ‘realms’ (Australasia, Afrotropics, Indo-Malaya, Neotropics and Oceania) and six tropical terrestrial biomes (tropical moist forest (i.e., ‘rainforest’), montane grassland, desert, mangrove, tropical dry forest and tropical grassland (i.e., ‘savanna’)), which are displayed in Appendix S1, Fig. S1.1a. The assignments for each species were cross-referenced with monographs and previous studies of *Pterocarpus* (Rojo, 1972, Saslis-Lagoudakis et al., 2011, Klitgård et al., in prep), and with the Plants Of the World Online database (http://powo.science.kew.org/). All traits (fruit morphology, realm assignment and biome assignment) are then visualised onto the *Pterocarpus* phylogenetic tree in Appendix S1, Fig. S1.1b.

We then compared the fit of trait-independent and trait-dependent biogeographical models outlined by Klaus and Matzke (2020) for biogeographical realms and biomes in BioGeoBEARS. For realms, we tested between three default models in BioGeoBEARS, all of which incorporate dispersal, extinction and range switching. These models were DEC (Ree & Smith, 2008), which additionally accounts for vicariance and ‘subset’ sympatric speciation, DIVALIKE (Ronquist, 1997), which also accounts for vicariance, and BAYAREALIKE (Landis et al., 2013), which additionally accounts for widespread sympatric speciation but not vicariance. For biomes, we assessed the fit of Markov-*k* (Mk) models (Lewis, P. O., 2001) (i.e., BAYAREALIKE a+, d=e=0 in BioGeoBEARS) to accommodate equal rates of character evolution, after Kriebel et al. (2019). Markov-*k* more accurately models the inheritance of ecological preferences in daughter species and avoids bias towards large ancestral ranges compared to models such as DEC. We also included four trait-dependent models in our comparisons. The first two models only parameterised trait switching *(‘Trait_1Rate’* and *‘Trait_2Rates’*). Then, for realm, a trait-dependent extension of the DEC model was used to include trait switching and multipliers on dispersal rate (*DEC_t12_t21_m2*), and for biome a trait-dependent implementation of the Markov-*k* model was used, which included the same trait switching and dispersal parameters (‘*Markov-k’_t12_t21_m2*).

The best-fit model was chosen using AICc and Akaike weights, from which ancestral ranges were estimated across *Pterocarpus*. Ancestral ranges were plotted onto the time-calibrated tree from MCMCTREE using the function *‘plot_BioGeoBears_results()’* for realm and biome independently. Finally, for biome, we counted the number of transitions between the most likely ancestral biomes at each node, and produced a matrix of biome shifts. This matrix was visualised as a transition plot using the *plotmat* function in the R package ‘diagram’ (Soetaert, 2012). Further information regarding biogeographical data collection and analysis is available in Appendix S1, Supplementary Methods.

## Results

### Phylogeny and divergence dating

A mean of 17.7% of reads were successfully assembled onto the *Angiosperms*-353 bait sequences, with a range from 6.9% to 30.5%. Gene recovery is shown as a heatmap in Appendix S1, Fig. S1.2. Of the 353 loci targeted by *Angiosperms*-353, we discarded 35 loci which were potentially paralogous and 15 which were assembled poorly or had over 50% missing data across taxa.

Our ASTRAL analysis indicated that most bipartitions were well supported with local posterior probabilities (LPP) >0.8, with 1 being full support (Fig. 1) and a quartet score of 0.68. Within this tree, most generic splits are well supported (LPP >0.9) and largely support the monophyly of *Pterocarpus* with the exception of one species, *Pterocarpus dubius* (formerly *Etaballia dubia). Pterocarpus dubius* was resolved in a clade containing *Inocarpus, Maraniona* and *Paramachaerium*, which was in a sister position to *Pterocarpus*. Within *Pterocarpus*, there were two major subclades, one containing species from the Neotropics (the neotropical clade) and one containing species from the Palaeotropics (the palaeotropical subclade). However, this split received a relatively low support (LPP=0.43, quartet score = 34.48%). The concatenated RAxML analysis of the same 303 loci (Appendix S1, Fig. S1.3) recovered a nearly identical topology to the ASTRAL species tree (Fig. 1), and was much better resolved, with nearly all nodes having bootstrap values >95. Moreover, the split between neotropical and palaeotropical *Pterocarpus* species received full support (BS=100) in the inference based on the concatenated dataset.

**Figure 1:**
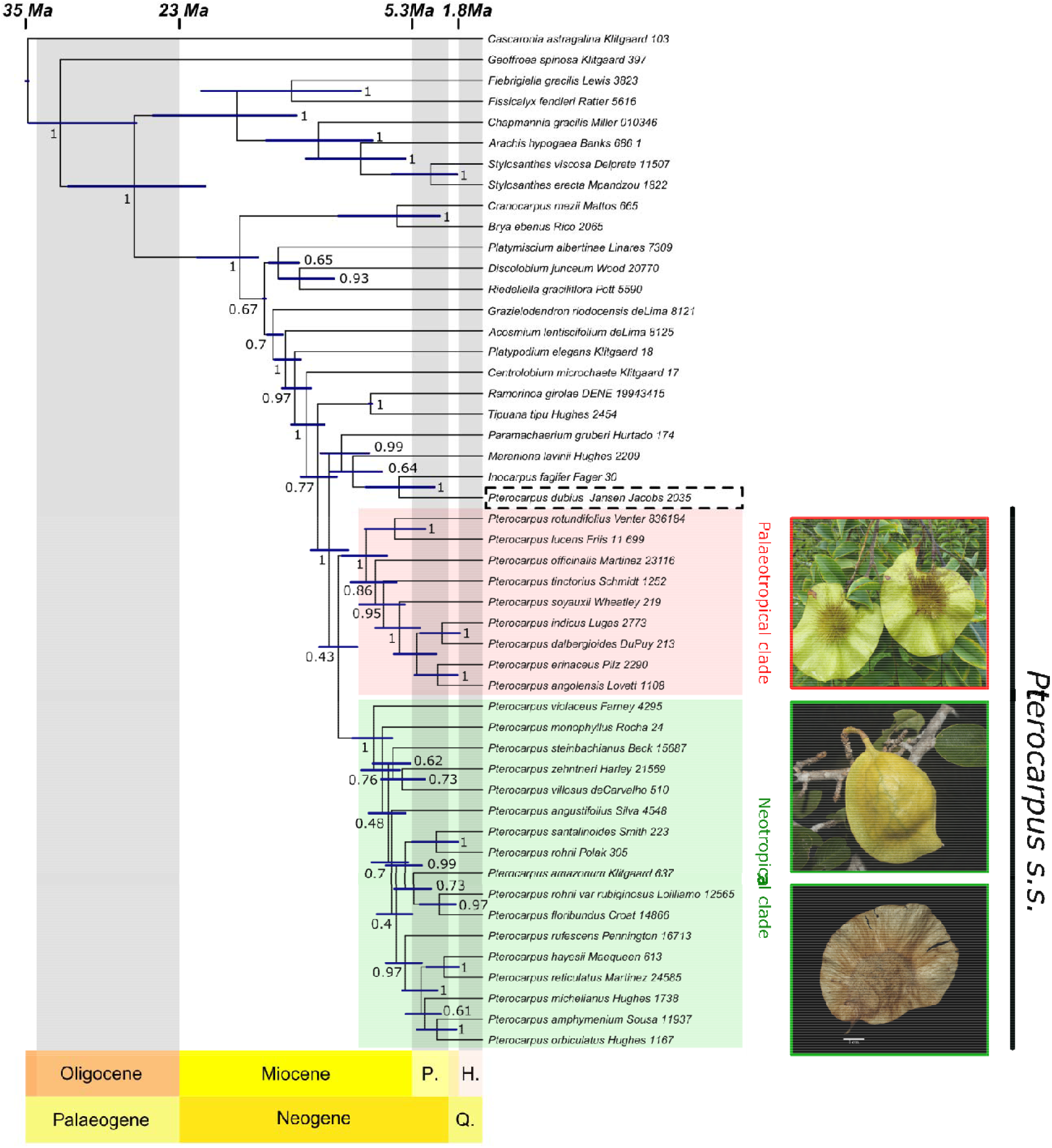
Fossil-calibrated MCMCtree of the Pterocarpus clade. Node bars show 95% highest posterior density (HPD) for node ages. Local posterior probabilities (LPP), a measure of topological support, are shown for each node, and were taken from the ASTRAL analysis which provided the tree topology for the MCMCtree analysis. Shading on the background of the phylogram represents geological epochs, which are labelled beneath the tree. The red and green shading indicates species belonging to the palaeotropical and neotropical subclades of *Pterocarpus*, respectively, and the extent of the genus *sensu stricto* is shown by the black bar. The dashed box around *Pterocarpus dubius* highlights its paraphyly with the rest of the genus. Photographs show winged fruits of *Pterocarpus erinaceus* (© Gwilym Lewis), coriaceous fruits of *Pterocarpus monophyllus* (*©* Domingos Cardoso) and winged fruits of *Pterocarpus violaceus* (© Reinaldo Aguilar, CC BY-NC-SA 2.0).

Divergence time estimation carried out in MCMCtree provided a robust, time-calibrated tree of the Pterocarpus clade. The two MCMC runs were convergent, with every node showing a suitable effective sample size (> 200), and posterior estimates fit the priors well as indicated in Appendix S1, Fig. S1.4a, b and c. The time-calibrated phylogenomic tree in Fig. 1 suggests that the Pterocarpus clade arose ~35 Ma, during the late Eocene, and diversification of present-day genera occurred during the Miocene, from 23-5.3 Ma. Of these, *Pterocarpus (sensu stricto)* diverged from its sister genera ~12 Ma, and diversified mostly in the late Miocene. It is worth noting that divergence date estimates are described by larger HPD intervals in the earlier divergence events within the Pterocarpus clade, and this increased uncertainty is mirrored in Appendix S1, Fig. S1.5. This figure indicates that uncertainty is greater in the earlier divergence events inferred, as well as being much reduced around fossil calibration points, as would be expected.

The topology of the BEAST trees (Appendix S1, Fig. S1.6) inferred using Sanger-sequencing data from GenBank recovered a broadly similar topology to the MCMCtree analysis (Fig. 1). Moreover, the divergence time estimates of this phylogeny were largely congruent with those from Fig. 1, including an origin of the Pterocarpus clade within the late Eocene (~38 Ma), followed by most divergence between genera, and diversification of *Pterocarpus*, within the Miocene (~23-5.3 Ma). However, the age estimate for the initial divergence of *Pterocarpus* is later than that inferred with MCMCtree, at around 25 Ma, again during the Miocene. In addition, the BEAST analysis displayed a much broader degree of uncertainty on age estimates, as indicated by the breadth of the 95% HPDs, and recovered *Pterocarpus dubius* as nested within *Pterocarpus*, whereas in the MCMCtree analysis *P. dubius* is in sister clade position to the rest of *Pterocarpus*.

### Biogeography

AICc comparison indicated that the best-fitting BioGeoBEARS model was DIVALIKE (AICc= 52.785) (Appendix S1, Table S1.5). Using DIVALIKE, we estimated a shared ancestral range of both the Neotropics and Afrotropics for the common ancestor of *Pterocarpus*, with subsequent splitting into neotropical and palaeotropical clades (Fig. 2a). The neotropical clade diversified only within the Americas and gave rise to the most species. Within the palaeotropical clade the common ancestor of *P. indicus* and *P. dalbergioides* appears to have dispersed into Asia, and subsequently into Australasia and Oceania, having a shared Afro-Asian range. It is interesting to note that within both neotropical and palaeotropical clades, there are species which share ranges, being found in both eastern South America and west Africa (*P. santalinoides* and *P. officinalis*). A shared characteristic of these two amphi-Atlantic species are thick-walled, wingless, buoyant fruits which allow water-borne dispersal. The current distribution of *Pterocarpus* species (both in terms of biome and realm), as well as the distribution of fruit dispersal traits across the ASTRAL tree, are visualised in Appendix S1, Fig. S1.1b.

**Figure 2a).**
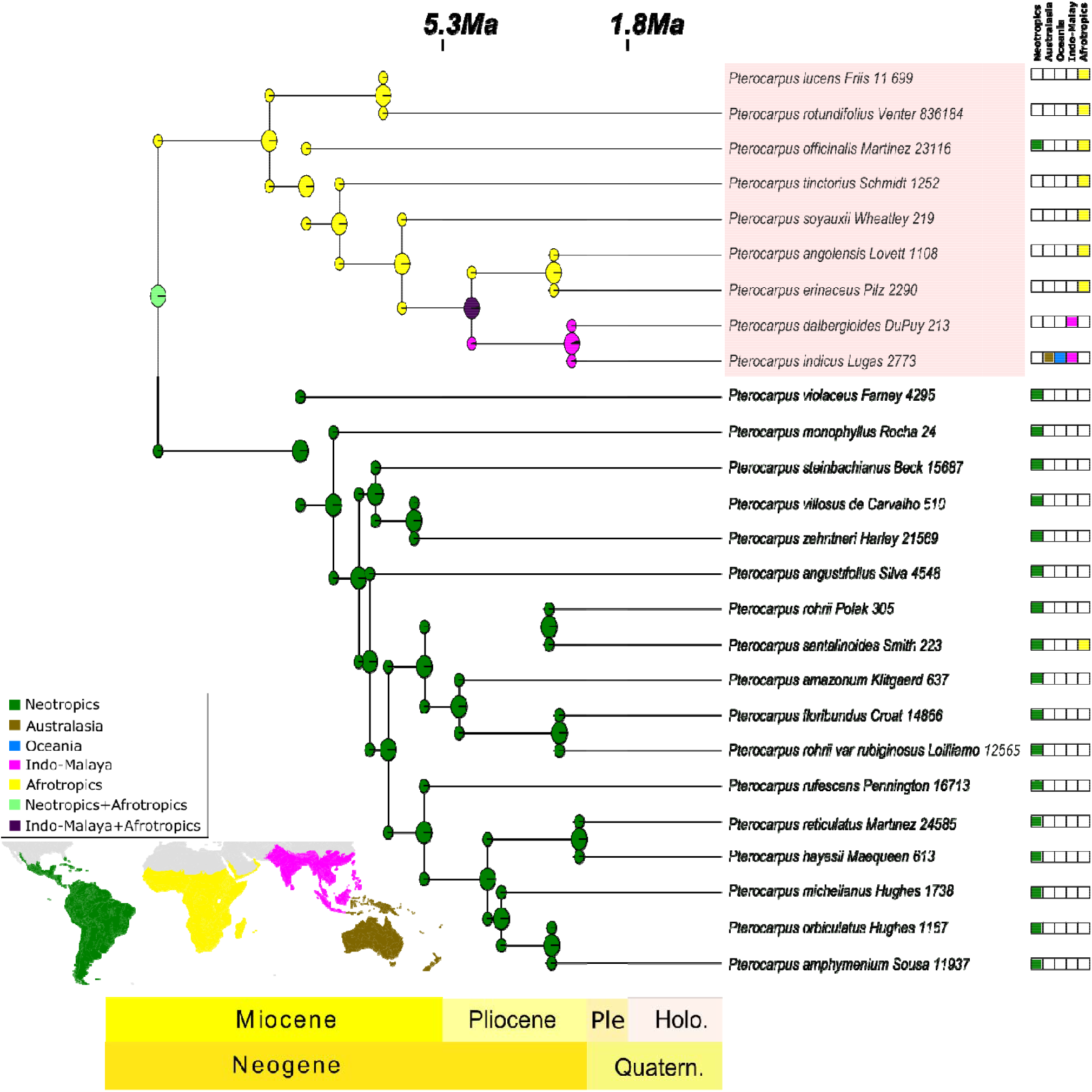
Biogeographical range estimation of *Pterocarpus*. Pie charts represent likelihoods of ranges derived from ancestral range estimations run on the MCMCtree topology, using the *R* package ‘BioGeoBEARS’. The legend represents five biogeographical realms defined by Olson *et al*. (2001) and shown in Appendix S1, Fig. S1.1a. Shading on the background of the phylogenetic tree represents geological epochs, and within the bottom bar ‘Ple’ represents ‘Pleistocene’, ‘Holo.’ represents ‘Holocene’ and ‘Quatern.’ represents ‘Quaternary’. Inset is a map displaying biogeographical realms as defined by Olson *et al*. (2001).

For biome evolution, the best-fit model was the trait-independent implementation of Markov-*k* (AICc= 328.218, Appendix S1, Table S1.5). Moist forest, dry forest and grassland were inferred as being nearly equally likely as the ancestral biome for the MRCA of *Pterocarpus* (Fig. 2b). Within the neotropical clade of *Pterocarpus*, there appears to be multiple polymorphic biome preferences, with species being found in dry forest as well as one other habitat type (desert, grassland or moist forest). However, for the majority of stem nodes within the neotropical *Pterocarpus* species, the most likely ancestral biome appears to have been dry forest, with only the clade containing *P. rohrii, P. santalinoides, P. amazonum* and *P. floribundus* containing species only found in moist forest. Within the palaeotropical *Pterocarpus* clade, the most probable state for the majority of ancestral nodes was shared between moist forest and grassland, with most species being found in grassland in addition to other biomes, mirroring the high polymorphism in neotropical *Pterocarpus* species. It is also of interest that one of the two *Pterocarpus* species which inhabit both South America and Africa (*P. officinalis*) occurs both in tropical rainforest as well as mangroves.

**Figure 2b).**
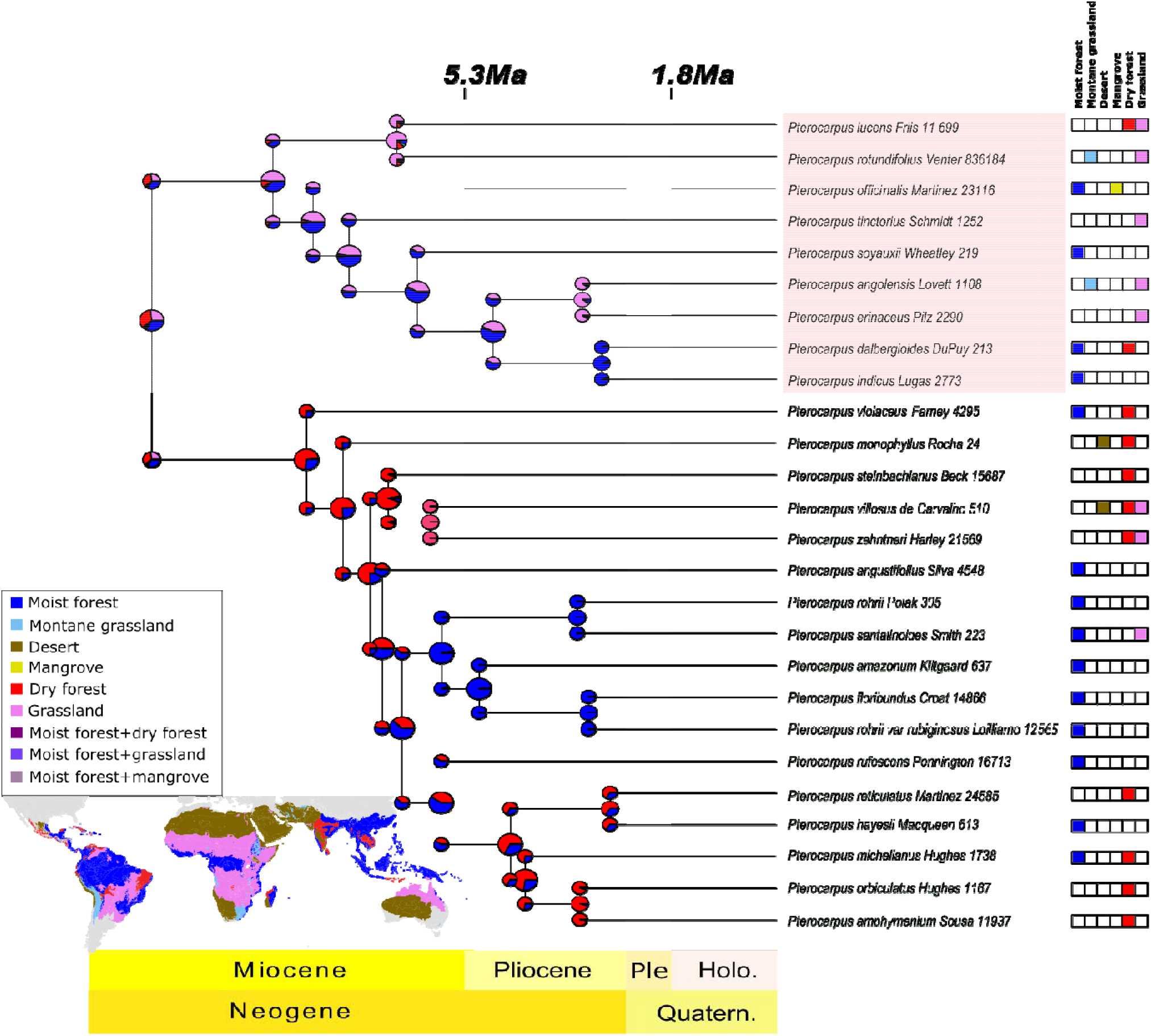
Historical biome estimation of *Pterocarpus*. Pie charts represent likelihoods of ranges derived from ancestral biome estimations run on the MCMCtree topology, using the *R* package ‘BioGeoBEARS’. The legend represents six biomes defined by Olson *et al*. (2001) and shown in Appendix S1, Fig. S1.1a. Shading on the background of the phylogenetic tree represents geological epochs, and within the bottom bar ‘Ple’ represents ‘Pleistocene’, ‘Holo.’ represents ‘Holocene’ and ‘Quatern.’ represents ‘Quaternary’. Inset is a map displaying biomes as defined by Olson *et al*. (2001).

We inferred the most shifts between biome states across the *Pterocarpus* tree into moist forests (i.e., rainforest) and grassland (i.e., savanna). Specifically, we recovered the most shifts from dry forest to moist forest (n=4, Fig. 3), followed by shifts from rainforest to grassland (n=3). Transitions also occurred between drier habitats (e.g. inferred shifts from dry forest to desert and grassland (n=2), or grassland to dry forest and desert (n=1)), between similar habitats (e.g., grassland into montane grassland (n=2), from moist forest to mangrove (n=1)) and in the contrasting direction to the most frequent shifts (e.g. moist forest to dry forest (n=2) and grassland to moist forest (n=1)).

**Figure 3).**
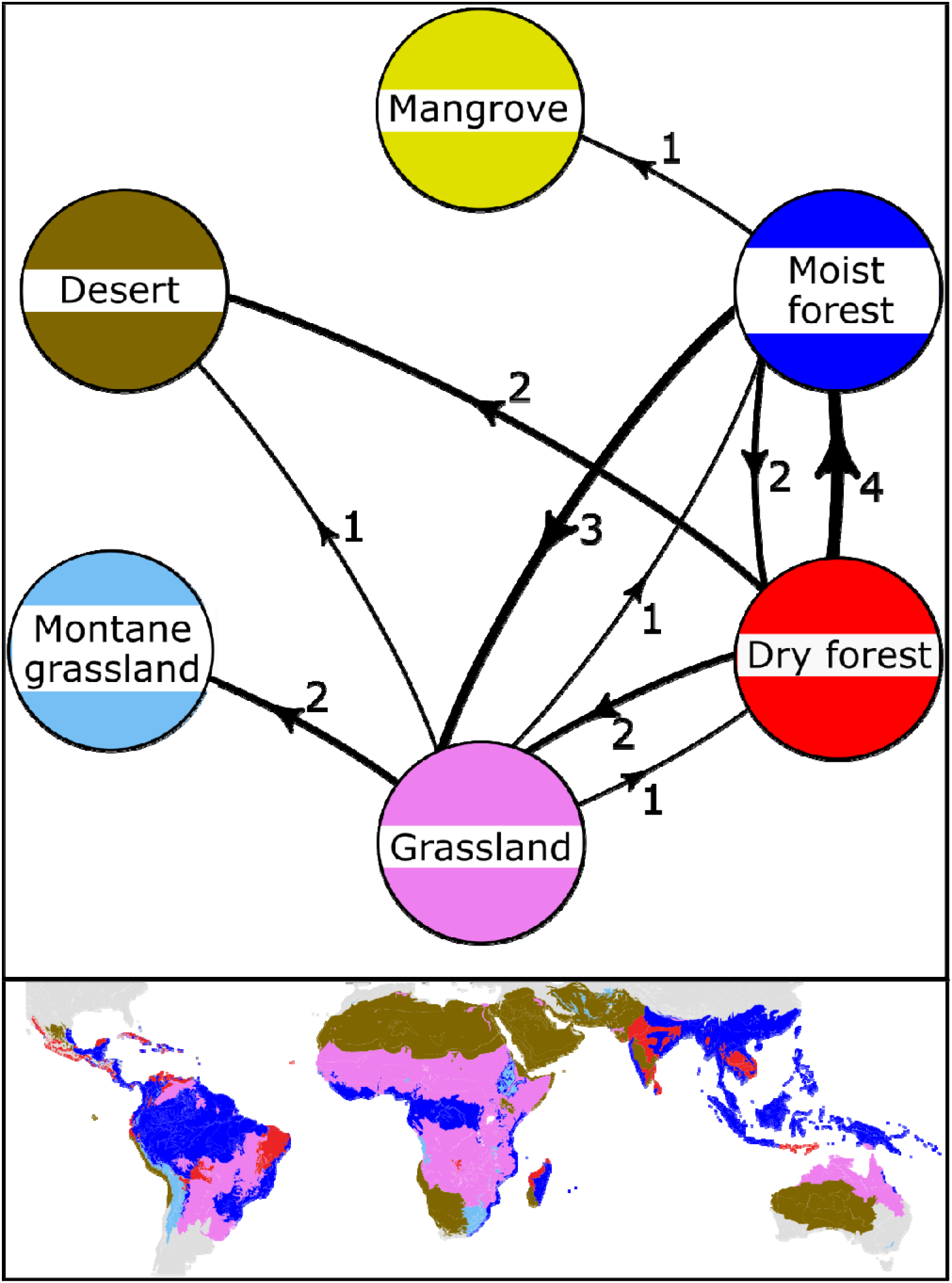
Biome transitions between species within *Pterocarpus*, based on the ancestral biome reconstruction with the M*k* model shown in Fig. 2b. Arrows on lines show the direction of the biome transition, while numbers and thickness of the lines represent the number of transitions between each biome. Inset is a map displaying biomes as defined by Olson *et al*. (2001).

## Discussion

Divergence dating indicated that the Pterocarpus clade began to diversify in the late Eocene (~35 Ma), followed by extensive diversification during the Miocene (23-5.3 Ma). During the Miocene, the common ancestor of *Pterocarpus* diverged into two clades. One clade was palaeotropical and diversified largely in Africa with subsequent dispersal into Indo-Malaya, Australasia and Oceania. The second clade was neotropical, which diversified only in the tropical Americas and was more speciose than the palaeotropical clade. The diversification of *Pterocarpus* appears to have involved multiple biome shifts, largely into lowland habitats (namely, savanna and tropical moist forest), and while dispersal across large geographical areas played a role in range expansion, the fruit dispersal traits we examined were not significantly associated with macroevolutionary dynamics according to our model comparisons.

Overall, it appears that traits for hydrochoric dispersal evolved multiple times independently in *Pterocarpus*, likely in concert with adaptation to waterlogged habitats such as mangroves. These traits which evolved to promote dispersal in the ecological theatre may have then led to rare long-distance dispersal across oceans (as we see in extant species such as *P. santalinoides* and *P. officinalis*). A similar pattern has been shown in the Bombacoideae (Malvaceae) (Zizka et al., 2020), as well as in *Centrolobium* and *Platymiscium*, which are close relatives of *Pterocarpus* (Klitgård, 2005, Pirie et al., 2009, Saslis Lagoudakis et al., 2008).

### Phylogenetic analysis of the Pterocarpus clade

Most Pterocarpus clade lineages, including *Pterocarpus* itself, arose during the Miocene, between 23-5.3 Ma (Fig. 1; Appendix S1, Fig. S1.6). This was congruent across methods (MCMCTREE, BEAST) and across data types (target capture data, Sanger-sequencing data). While our divergence dating analyses were broadly congruent with each other across methods and data type, they differed in their date estimates for the origin of *Pterocarpus*. MCMCTREE analysis inferred an origin date of 12Ma (Fig. 1), whereas the BEAST analysis suggested that *Pterocarpus* arose around 25Ma (Appendix S1, Fig. S1.6). The most likely reason for this discrepancy is the size of the dataset, since MCMCtree was run on a nuclear target capture dataset comprising 303 loci, while BEAST was run on pre-existing Sanger sequence data for 5 loci, resulting in a much less informative sequence matrix.

It is also likely that the inclusion of plastid loci within the BEAST analysis extended the date estimate for the origin of *Pterocarpus*, due to deeper divergences within the plastid genome (Wolfe et al., 1987). This conflicting signal would have also led to the much higher degree of uncertainty on age estimates in the BEAST analysis, as indicated by the breadth of the 95% HPDs. In addition, the nested position of *Pterocarpus dubius* within *Pterocarpus*, as shown in the BEAST analysis (Appendix S1, Fig. S1.6) contrasts with the sister clade position of *P. dubius* to *Pterocarpus* in the MCMCTREE analysis (Fig. 1). This species was previously circumscribed as *Etaballia dubia* and is highly divergent in terms of floral morphology, however previous work with Sanger-sequencing loci resolved this species within *Pterocarpus* (Klitgård et al., 2013). It is possible that the inclusion of chloroplast loci in the BEAST tree caused this incongruence, due to the lower mutation rate of chloroplast regions (Wolfe et al., 1987), or through plastid capture resulting from hybridisation (Naciri & Linder, 2015). While there is undoubtedly useful signal within these plastid loci, further work using plastomes would help to accurately disentangle these discordances.

### *Spatio-temporal evolution of* Pterocarpus

The Miocene-era diversification of the Pterocarpus clade, and particularly *Pterocarpus*, reflects patterns shown in other tropical tree genera (e.g., the mahoganies (Meliaceae) (Koenen et al., 2015), the Brownea clade (Leguminosae) (Schley et al., 2018) and the Daniellia clade (Leguminosae) (Choo et al., 2020)). This diversification was likely concurrent with major climatic, geological and ecological changes which occurred in the tropics during the Miocene (Keeley & Rundel, 2005, Morley, 2000) such as was the Miocene global cooling event (Zachos et al., 2001, 2008). This cooling was precipitated by the closure of the Tethys seaway and a subsequent change in ocean currents (Rommerskirchen et al., 2011, Zhang, Z. et al., 2014), leading to a decrease in global temperature, desiccation of the continents and a retreat of tropical vegetation, especially on the African continent (Couvreur et al., 2021, Morley & Richards, 1993, Senut et al., 2009). Such cooling likely also explains the long-distance dispersal events and biome-switching we observed in our ancestral range reconstructions for *Pterocarpus* (Fig. 2a & 2b), particularly within the palaeotropical clade, as discussed below.

#### i The palaeotropical clade

The palaeotropical clade appears to have diversified largely within Africa over the last ~10Ma, in the late Miocene and into the early Pliocene (Fig. 2a), with multiple biome shifts from moist forests into grasslands (n=3, Fig. 3). These events were likely concurrent with the cooling and desiccation of the African continent which lead to the expansion of grasslands during this period (Couvreur et al., 2021). This palaeotropical clade was also less speciose than its neotropical counterpart, which may be explained by higher levels of extinction in rainforest lineages, again precipitated by the expansion of grassland in Africa’s past. This has led to the well-documented ‘odd man out’ pattern of rainforest tree diversity, with the Afrotropics being significantly less diverse that other tropical regions (Couvreur, 2015, Richards, 1973). Moreover, most of the extant *Pterocarpus* species within the palaeotropical clade are now found in grassland and are wind-dispersed (Fig. 2b; Appendix S1, Fig. S1.1b), suggesting historical adaptation to novel grassland environments.

Within the Palaeotropical clade, the two Asian sister species (*P. dalbergioides* and *P. indicus)* only dispersed to the Asian tropics and beyond into Australasia and Oceania around 4Ma, during the Pliocene (Fig. 2a). It is likely that the divergence of these species was caused by infrequent eastward seed dispersal, promoted by the east African coastal current (EACC) between Africa and Asia which formed with the closing of the Tethys seaway in the late Miocene (Scotese, 2004). It is interesting to note that these species are found in rainforests, as rainforest tree species tend to be excellent dispersers (Dexter et al., 2017, Pennington & Lavin, 2016), and are often water dispersed. This mirrors the highly dispersible nature of the other rainforest-dwelling member of the Palaeotropical clade (*P. officinalis*) which is distributed in both Africa and South America and is also found in mangroves. Moreover, *P. officinalis* is derived from Miocene African ancestors (Fig. 2a), is present in the Neotropical fossil record (Graham, 1995) and includes a West African subspecies (subsp. g*illetii*), further suggesting that the corky exocarp of the fruit, which it evolved for dispersal at an ecological scale (Appendix S1, Fig. S1.1b), is responsible for rare dispersal across the Atlantic and the foundation of a Neotropical population (Rivera Ocasio et al., 2002).

#### ii The neotropical subclade

Within neotropical *Pterocarpus* we inferred multiple events of biome-switching from dry forest into rainforest (Fig. 2b; n=4, Fig. 3)), with the most likely ancestral biome for this clade being dry forest. These events mainly occurred between the late Miocene and the Pliocene (~8 - 2 Ma). In particular, we see the switch from dry forest in stem nodes into clades which are found nearly exclusively in wet forest towards the tips, mirroring findings in closely related genera (e.g. *Platymiscium* (Saslis Lagoudakis et al., 2008)). Indeed, biome switching between dry forest and moist forest, which are often interdigitated, is well documented in the Neotropics (Dexter et al., 2018, Ireland et al., 2010, Klitgård, 2005, Pennington & Dick, 2010, Pezzini et al., 2020).

The majority of Neotropical dry forests formed during the Miocene (Pennington & Ratter, 2006) as shown by fossil evidence (e.g. Burnham, 1995, Burnham & Carranco, 2004). This was followed by periods of intense climatic flux and the contraction of dry forests resulting from increasingly wet Pleistocene interglacial periods, leading to the ‘refugia’ of dry forest which we observe today within inter-Andean valleys, Central America and the ‘dry arc’ south of the Amazon in Brazil and Bolivia (Pennington et al., 2004). The biome switching we inferred within the Neotropics may be explained by these oscillations, as the expansion of rainforests would have resulted in a much higher likelihood of immigration into rainforest by plant species dispersing from dry forest refugia.

#### iii *Asymmetrical biome transitions in* Pterocarpus

Biome transitions across *Pterocarpus* occurred mainly into moist forest (i.e., rainforest) and grassland (i.e., savanna) (Fig. 3). The higher relative frequency of immigration into both moist forests and grasslands may be explained by the fact that these communities experience persistent disturbance (e.g., fire in tropical savannas) or because they contain species which are vulnerable to climatic perturbation (e.g., drought in rainforests) (da Costa et al., 2010, Pennington & Lavin, 2016). In rainforests, intermediate disturbance increases the likelihood of seedling establishment across entire communities due to intense competition for light (Hubbell et al., 1999), and within savannas, seedling establishment of local species is greatly reduced by frequent fire (Hoffmann, 1996, Setterfield, 2002, Wakeling et al., 2011). This may promote the competitive success of tree seeds dispersing into such environments from adjacent habitats. Thus, the highly dispersible nature of *Pterocarpus* fruits could have promoted immigration into these novel biomes, and the moderate disturbance which occurs to plant communities within these biomes could have facilitated establishment, adaptation and speciation.

The frequency of biome shifts into savanna and moist forests also implies that their wide extent renders them more likely to exchange seeds with adjacent biomes (Donoghue & Edwards 2014). This is particularly true for the ‘donation’ of immigrants from restricted biomes which interdigitate with savannas or rainforest, such as dry forest. Overall, our results highlight that biome switching may play an important role in diversification of tropical trees and is more common than previously thought, as recently highlighted across neotropical taxa (Antonelli et al., 2018).

### Caveats

Despite the utility of biogeographical models for understanding spatio-temporal evolution, they are not without their drawbacks. In particular, biogeographical models may be prone to over-estimating long-distance dispersal when model assumptions are violated (e.g., *DEC+J* in BioGeoBEARS (Ree & Sanmartín, 2018)), highlighting the importance of appropriate model choice (Matzke, 2014). As far as is possible, we have attempted to account for these biases by, for example, avoiding the use of the *+J* parameter in biogeographical analyses.

It is also possible that small sample sizes may be problematic, since with larger datasets parameter estimates are likely to be more accurate (e.g., in species distribution modelling (Wisz et al., 2008)). However, as in our study, this can be avoided by having a balanced sampling strategy which accounts for the nearly all species in studies to <50 taxa (e.g., in Araliaceae (Valcárcel & Wen, 2019), penguins (Vianna et al., 2020) and cat-eyed snakes (Weinell et al., 2021)). Finally, our specimens were identified by taxonomic experts on *Pterocarpus* and underwent multiple corroborative phylogenetic analyses, thereby minimising error as a result of misidentification or phylogenetic uncertainty. However, even while accounting for such biases, it is important to interpret biogeographical model outputs cautiously, and to realise that they are approximate estimates of biogeographic history.

## Conclusion

Overall, our analyses suggest that while a dispersal trait-dependent model of range macroevolution was not supported, the biogeographical history of *Pterocarpus* was likely mediated by cross-continental dispersal, associated range change and diversification into adjacent biomes. Indeed, biome switching between moist and arid environments, likely precipitated by infrequent long-distance dispersal events and environmental change, appear to have promoted ecological differentiation and diversification within *Pterocarpus* since the early Miocene.

## Acknowledgements

All the authors would like to thank the PAFTOL program at RBG, Kew for funding the sequencing of Pterocarpus clade accessions. MQ’s lab visit was funded by the National Natural Science Foundation of China (grant no. 31670193) and the Science and Technology Planning Project of Guangdong Province (grant no. 2019B030316020).

## Data availability statement

The target capture sequencing data generated for this project are publicly available in the NCBI sequence read archive under the BioProject number PRJNA728569, with the accession numbers SAMN19092119 - SAMN19092167.

## Significance statement

Dispersal involves organisms spreading from one location to another, and is one of the major processes defining the distribution of organisms. Sometimes, structures which evolved for local dispersal may lead to rare, long-distance dispersal, and this may allow species to colonise new landmasses and diversify in novel biomes. As such, here we build an evolutionary tree to understand relationships between a genus of rosewood species (*Pterocarpus*), and test whether dispersal traits and biome switching have impacted their evolution. We found that dispersal and biome switching, likely precipitated by climate change in the Miocene, impacted the modern distribution of Pterocarpus, but that this was not significantly explained by dispersal structures.

## Biosketch

Rowan Schley is a postdoctoral researcher interested in the evolution, biogeography and conservation of tropical biodiversity. His research is broadly centred around speciation with a particular focus on the effects of introgression, using phylogenomic and population genomic approaches to address these questions in tropical trees.

R.J.S. conceived the hypotheses, led data analysis and wrote the manuscript. M.Q. and P.M. led the phylogenomic data generation. M.V., M.d.l.E. and G.L. provided many comments and background information on the Legume family for the manuscript. B.K. helped conceive the study and led sample collection, and provided many comments on the manuscript. All authors contributed to editing the manuscript.

