## Appendix S1 for "The diversification of *Pterocarpus* (Leguminosae: Papilionoideae) was influenced by biome-switching and infrequent long-distance dispersal"

**Appendix S1** of Supporting Information for:

**The diversification of *Pterocarpus* (Leguminosae: Papilionoideae) was influenced by biome-switching and infrequent long-distance dispersal**

Rowan J. Schley, Ming Qin, Mohammad Vatanparast, Panagiota Malakasi, Manuel de la Estrella, Gwilym Lewis, Bente Klitgård

**Tables**

| **Species** | **Voucher** | **Collection locality** |
| --- | --- | --- |
| *Acosmium lentiscifolium* Schott * | de Lima 8125 (K) | Brazil |
| *Arachis hypogaea* L. * | Banks 686.1 (K) | Ecuador |
| *Brya ebenus* DC. * | Rico 2065 (K) | Cuba |
| *Cascaronia astragalina* Griseb. * | Klitgård & Lewis 103 (K) | Argentina |
| *Centrolobium microchaete* (Mart. ex Benth.) H.C. Lima * | Klitgård 17 (NHM) | Brazil |
| *Chapmannia gracilis* (Balf. f.) Thulin * | Miller 0.10346 (K) | Yemen |
| *Cranocarpus mezii* Taub. * | Mattos 665 (K) | Brazil |
| *Discolobium junceum* Micheli * | Wood 20770 (K) | Bolivia |
| *Fiebrigiella gracilis* Harms * | Lewis 3823 (K) | Ecuador |
| *Fissicalyx fendleri* Benth. * | Ratter 5616 (K) | Brazil |
| *Geoffroea spinosa* Jacq. * | Klitgård 397 (NHM) | Ecuador |
| *Grazielodendron riodocensis* H.C.Lima * | de Lima 8121 (K) | Brazil |
| *Inocarpus fagifer* (Parkinson) Fosberg * | Fager & Junaai 30 (NHM) | Singapore Botanical Garden (cultivated) |
| *Maraniona lavinii* C.E.Hughes, G.P.Lewis, Daza & Reynel * | Hughes 2209 (K) | Peru |
| *Paramachaerium gruberi* Brizicky * | Hurtado 174 (K) | Costa Rica |
| *Platymiscium albertinae* Standl. & L.O. Williams * | Linares 7309 (K) | Honduras |
| *Platypodium elegans* Vogel * | Klitgård 18 (NHM) | Brazil |
| *Pterocarpus amazonum* (Mart. ex Benth.) Amshoff | Klitgård 637 (NHM) | Ecuador |
| *Pterocarpus amphymenium* DC. | Sousa 11937 (K) | Mexico |
| *Pterocarpus angolensis* DC. | Lovett & Congden 1108 (K) | Tanzania |
| *Pterocarpus angustifolius* (Benth.) Klitg. & Mansfield-Williams | Silva 4548 (K) | Brazil |
| *Pterocarpus dalbergioides* Roxb. | Du Puy 213 (K) | Madagascar |
| *Pterocarpus dubius* (Kunth.) Spreng. | Jansen-Jacobs 2035 (K) | Guyana |
| *Pterocarpus erinaceus* Poir. | Pilz 2290 (K) | Nigeria |
| *Pterocarpus floribundus* Pittier | Croat 14866 (MO) | Panama |
| *Pterocarpus hayesii* Hemsl. | Macqueen 613 (K) | Costa Rica |
| *Pterocarpus indicus* Willd. | Lugas 2773 (K) | Malaysia |
| *Pterocarpus lucens* Lepr. ex Guill. & Perr. | Friis 11.699 (K) | Ethiopia |
| *Pterocarpus michelianus* N.Zamora | Hughes 1738 (K) | Costa Rica |
| *Pterocarpus monophyllus* Klitg., L.P.Queiroz & G.P.Lewis | Rocha 24 (K) | Brazil |
| *Pterocarpus officinalis* Jacq. | Martinez 23116 (K) | Guatemala |
| *Pterocarpus orbiculatus* DC. | Hughes 1167 (K) | Mexico |
| *Pterocarpus reticulatus* Standl. | Martinez 24585 (MO) | Mexico |
| *Pterocarpus rohrii* Vahl | Polak 305 (K) | Guyana |
| *Pterocarpus rohrii* var. *rubiginosus* Schery | Loilliamo 12565 (K) | - |
| *Pterocarpus rotundifolius* (Sond.) Druce | Venter 836184 (K) | Botswana |
| *Pterocarpus rufescens* Benth. | Pennington 16713 (K) | Peru |
| *Pterocarpus santalinoides* L'Hér. ex DC. | Smith 223 (K) | Peru |
| *Pterocarpus soyauxii* Taub. | Wheatley 219 (K) | Cameroon |
| *Pterocarpus steinbachianus* Harms | Beck 15687 (K) | Bolivia |
| *Pterocarpus tinctorius* Welw. | Schmidt 1252 (K) | Tanzania |
| *Pterocarpus villosus* Mart.ex Benth. | de Carvalho 510 (K) | Brazil |
| *Pterocarpus violaceus* Vogel | Farney 4295 (K) | Brazil |
| *Pterocarpus zehntneri* Harms | Harley 21569 (K) | Brazil |
| *Ramorinoa girolae* Speg. * | DENE? 1994-3415 (K) | Argentina |
| *Riedeliella graciliflora* Harms * | Pott 5590 (K) | Brazil |
| *Stylosanthes erecta* P.Beauv * | Mpandzou 1822 (K) | Congo Brazzaville |
| *Stylosanthes viscosa* (L.) Sw. * | Delprete 11507 (K) | French Guiana |
| *Tipuana tipu* (Benth.) Kuntze * | Hughes 2454 (K) | Bolivia |

**Table S1.1:** Sampled vouchers, collector information and collection localities for *Pterocarpus* species used in phylogenomics, MCMCtree and biogeographical inference. Herbaria from which accessions were collected are cited next to the collector name and number. Outgroup taxa from the Pterocarpus clade are marked with an asterisk (*).

| **Cycle step** | **Temperature** | **Time** | **N^o^ Cycles** |
| --- | --- | --- | --- |
| *Initial Denaturation* | 98°C | 30 seconds | 1 |
| *Denaturation* | 98°C | 10 seconds | 12 |
| *Annealing/Extension* | 65°C | 75 seconds |  |
| *Final Extension* | 65°C | 5 minutes | 1 |
| *Hold* | 4°C | ∞ | - |

**Table S1.2:** Amplification conditions used in the NEBNext ® Ultra™ II protocol to amplify adaptor-ligated DNA libraries.

| **Taxon** | **Collector** | **Collection locality** | **ndhF-rpl32** | **rbcl** | **matK** | **nrITS** | **trnL-trnF** |
| --- | --- | --- | --- | --- | --- | --- | --- |
| *Centrolobium microchaete*  (Mart. ex Benth.) H.C. Lima* | Klitgård 17 (AAU) | Brazil,  Minas Gerais | EU735853 | JN083700 | EU401408 | JN083771 | EU735859 |
| *Grazielodendron riodocensis* H.C.Lima * | Klitgård 23 (AAU) | Brazil,  Espírito Santo | EU735854 | JN083701 | AF270862 | JN083772 | EU735864 |
| *Inocarpus fagifer*  (Parkinson) Fosberg * | Fager & Junaai 30 (MT) | Singapore Botanic Garden (cult.) | JN083707 | JN083702 | AF270878 | JN083773 | EU735865 |
| *Maraniona lavinii*  C.E.Hughes, G.P.Lewis, Daza & Reynel * | Hughes 2209 (K) | Peru, Cajamarca, Balsas | KF436449 | KF436463 | KF436439 | KF436422 | KF436480 |
| *Maraniona lavinii*  C.E.Hughes, G.P.Lewis, Daza & Reynel * | Hughes 2647 (K) | Peru, Amazonas | KF436450 | KF436464 | KF436440 | KF436423 | KF436481 |
| *Paramachaerium ormosioides* (Ducke) Ducke * | Sabatier & Molino 5206 (K) | French Guiana, Inselbergs de la Haute Wanapi | – | – | KF436441 | KF436427 | KF436485 |
| *Platymiscium pubescens*  subsp. *fragrans* (Rusby) Klitgård * | Nee 37050 (K) | Bolivia, Santa Cruz | EU735910 | JN083704 | EU735968 | - | - |
| *Pterocarpus acapulcensis*  Rose | Hughes 769 (K) | Venezuela, Trujillo | JN083584 | JN083708 | JN083532 | JN083464 | JN083638 |
| *Pterocarpus acapulcensis*  Rose | Wurdack 41202 (K) | Venezuela, Bolivar | JN083585 | JN083709 | JN083533 | JN083465 | JN083639 |
| *Pterocarpus amazonum*  (Mart. ex Benth.) Amshoff | Klitgård 637 (BM) | Ecuador, Napo | JN083587 | JN083711 | JN083535 | JN083467 | JN083641 |
| *Pterocarpus amazonum*  (Mart. ex Benth.) Amshoff | Schunke 6192 (K) | Peru, Huanuco | JN083586 | JN083710 | JN083534 | JN083466 | JN083640 |
| *Pterocarpus amphymenium*  DC. | Hughes 1167 (K) | Mexico, Guerrero | JN083591 | JN083715 | JN083539 | JN083471 | JN083645 |
| *Pterocarpus amphymenium*  DC. | Medrano 11805 (K) | Mexico, Oaxaca | JN083588 | JN083712 | JN083536 | JN083468 | JN083642 |
| *Pterocarpus amphymenium*  DC. | Morton & Makrinius 2399 (K) | Mexico, Oaxaca | JN083590 | JN083714 | JN083538 | JN083470 | JN083644 |
| *Pterocarpus amphymenium*  DC. | Sousa 11937 (K) | Mexico, Oaxaca | JN083589 | JN083713 | JN083537 | JN083469 | JN083643 |
| *Pterocarpus angolensis*  DC. | Rodin 8929 (K) | Namibia, Oshikango | JN083593 | JN083717 | – | JN083473 | JN083646 |
| *Pterocarpus brenanii*  Barbosa & Torre | Cannell 16 (K) | Zimbabwe (Rhodesia), Distr. Kariba, (Sanyati river) | JN083594 | JN083718 | JN083540 | JN083475 | JN083647 |
| *Pterocarpus cf. erinaceus*  Poir. | Chapman 5213 (K) | Nigeria, Gongola State, Sarduana L.G. area, Mambilla Plateau, Akwaijantar forest | JN083595 | JN083719 | JN083541 | JN083476 | JN083648 |
| *Pterocarpus cf. reticulatus*  Standl. | Monro 3646 (BM) | El Salvador, La Libertad | KF436456 | KF436471 | KF436442 | – | – |
| *Pterocarpus cf. reticulatus*  Standl. | Ireland 1 (K) | Mexico, Chiapas | – | KF436470 | – | KF436430 | KF436488 |
| *Pterocarpus dalbergioides*  Roxb. | Du Puy 213 (K) | Madagascar, Toamasina (Tamatave Prov.), NE of Maroansetta | JN083596 | JN083720 | – | JN083477 | JN083649 |
| *Pterocarpus dubius*  (Kunth) Spreng. | Chanderbali 210 (K) | Guyana,  Takutu, Upper Essequibo | – | KF436459 | KF436437 | KF436420 | - |
| *Pterocarpus erinaceus*  Poir. | Pilz 2290 (K) | Nigeria, Ogun | JN083597 | JN083721 | JN083542 | JN083478 | JN083650 |
| *Pterocarpus erinaceus*  Poir. | Velakamp 6117 (K) | Ghana, Amezofe | JN083598 | JN083722 | JN083543 | JN083479 | JN083651 |
| *Pterocarpus floribundus*  (Benth.) Kuntze | Croat 14866 (K) | Panama, Canal Zone, Barro Colorado Island | – | KF436472 | – | KF436431 | KF436489 |
| *Pterocarpus hayesii*  Hemsl. | Monro 3668 (BM) | El Salvador, La Libertad, | – | KF436473 | – | KF436432 | KF436490 |
| *Pterocarpus indicus*  Willd. | Ambri & Arifin AA401 (K) | Indonesia,Wanariset, Kalimantan Timur | JN083599 | JN083724 | JN083545 | JN083481 | – |
| *Pterocarpus indicus*  Willd. | Lugas 2773 (K) | Malaysia, Borneo, Kota Belud district | JN083600 | JN083725 | JN083546 | JN083482 | JN083653 |
| *Pterocarpus indicus*  Willd. | Stancik & Nathaniel 5198 (K) | Papua New Guinea, Madang, Baitabag | – | JN083723 | JN083544 | JN083480 | JN083652 |
| *Pterocarpus lucens* subsp. *antunesii*  (Taub.) Rojo | Pereira & Correia 2121 (K) | Mozambique (Mocambique), Cabora Bassa | JN083601 | JN083726 | JN083547 | JN083483 | JN083654 |
| *Pterocarpus lucens* Lepr. ex Guill. & Perr. subsp. *lucens* | Breteler 1155 (K) | Cameroon, Savannah, Money,Betare Oya | JN083603 | JN083728 | JN083549 | JN083485 | JN083656 |
| *Pterocarpus lucens* Lepr. ex Guill. & Perr. subsp*. lucens* | Diallo 295 (K) | Mali, (French Sudan), Azzanturi, Ilahatan | JN083604 | JN083729 | JN083550 | JN083486 | JN083657 |
| *Pterocarpus lucens* Lepr. ex Guill. & Perr. subsp*. lucens* | Demissew 1925 (K) | Ethiopia, Gojam Adm. Region, Metekel Awraja | JN083602 | JN083727 | JN083548 | JN083484 | JN083655 |
| *Pterocarpus macrocarpus*  Kurz | Monyrak & Meng 223 (K) | Cambodia, Prov. Stung Treng, Distr., Thala Barevath, Kalay Island Ramsar site | JN083605 | JN083730 | JN083551 | JN083487 | JN083658 |
| *Pterocarpus magnicarpus*  Schery | Daza 1376 (K) | Peru, San Ramon | – | KF436474 | KF436443 | – | KF436491 |
| *Pterocarpus marsupium*  Roxb. | Klackenberg & Lundin 253 (K) | India, Tamil Nadu, Nilgiris, Ootacamund area | – | JN083731 | – | JN083488 | JN083659 |
| *Pterocarpus michelianus*  N. Zamora | V. Ramirez 263 (MO) | Costa Rica, San Jose | – | JN083734 | JN083554 | JN083491 | JN083662 |
| *Pterocarpus mildbraedii* Harms  subsp. *mildbraedii* | Chapman 3849 (K) | Nigeria, NE State, Sardauna, River Nwam Forest Reserve, Mambilla Plateau | JN083608 | JN083735 | JN083555 | JN083492 | JN083663 |
| *Pterocarpus mildbraedii*  subsp. *usambarensis*  (Verdc.) Polhill | Greenway 7923 (K) | Tanzania, E of Usambara | JN083609 | JN083736 | – | JN083493 | JN083664 |
| *Pterocarpus monophyllus*  Klitgård, L.P.Queiroz & G.P.Lewis | Rocha 23 (K) | Brazil, Bahia | – | – | JN083556 | JN083494 | JN083665 |
| *Pterocarpus officinalis* Jacq. subsp. *officinalis* | Martinez 23116 (K) | Guatemala | JN083610 | JN083737 | JN083557 | JN083496 | JN083666 |
| *Pterocarpus officinalis* Jacq. subsp. *officinalis* | Boom 7054 (K) | Puerto Rico | JN083611 | JN083738 | JN083558 | JN083497 | JN083667 |
| *Pterocarpus osun*  Craib | Lowe 4702 (K) | Nigeria, Ibadan | – | JN083739 | – | JN083498 | JN083668 |
| *Pterocarpus rohrii*  Vahl | Elias 1209 (K) | Colombia, Barranquilla | JN083615 | JN083745 | JN083562 | JN083504 | JN083674 |
| *Pterocarpus rohrii*  Vahl | Hughes 1190 (K) | Guatemala | JN083613 | JN083741 | JN083560 | JN083500 | JN083670 |
| *Pterocarpus rohrii*  Vahl | Mori & Smith 25173 (K) | French Guiana | – | JN083742 | JN083561 | JN083501 | JN083671 |
| *Pterocarpus rohrii*  Vahl | Nee 50163 (K) | Bolivia, Santa Cruz | JN083616 | JN083746 | JN083563 | JN083505 | JN083675 |
| *Pterocarpus rohrii*  Vahl | Pirani 26889 (K) | Brazil,  Espírito Santo | JN083617 | JN083747 | JN083564 | JN083506 | JN083676 |
| *Pterocarpus rohrii*  Vahl | Tenorio 19712 (K) | Mexico, Chiapas | JN083612 | JN083740 | JN083559 | JN083499 | JN083669 |
| *Pterocarpus rotundifolius* subsp. *polyanthus*  (Harms) Mend. & Sousa | Bingham & Jeffery 11799 (K) | Zambia, S Prov., Mazabuka Distr., Kaleya Ranch | JN083618 | JN083748 | JN083565 | JN083507 | JN083677 |
| *Pterocarpus rotundifolius* subsp. *polyanthus* var*. martinii*  (Dunkley) Mend. & Sousa | Leach 14991 (K) | Zimbabwe (Rhodesia), Distr. Lomagundi | JN083620 | JN083750 | JN083567 | JN083509 | JN083679 |
| *Pterocarpus rotundifolius* subsp. *polyanthus* var*. martinii*  (Dunkley) Mend. & Sousa | White 6492 (K) | Zambia, S Prov., Mazabuka | JN083619 | JN083749 | JN083566 | JN083508 | JN083678 |
| *Pterocarpus rotundifolius* (Sond.) Druce  subsp. *rotundifolius* | Balsinhas 2841 (K) | South Africa, Transvaal, Pretoria Bot. Res. Inst., Botanic Gardens, Kloof Forest Section | JN083621 | JN083751 | JN083568 | JN083510 | JN083680 |
| *Pterocarpus santalinoides*  L'Hér. ex DC. | Hatschbach 65625 (K) | Brazil, Mato Grosso | JN083625 | JN083756 | JN083571 | JN083515 | JN083685 |
| *Pterocarpus santalinoides*  L'Hér. ex DC. | Lock 84/29 (K) | Togo, banks of River Koumangou at Naboulgou, Sansanne Mango | JN083624 | JN083755 | JN083570 | JN083514 | JN083684 |
| *Pterocarpus santalinoides*  L'Hér. ex DC. | Martin SL871 (K) | Sierra Leone, Taiama | JN083623 | JN083754 | JN083569 | JN083513 | JN083683 |
| *Pterocarpus santalinoides*  L'Hér. ex DC. | Tutin s.n. (K) | Senegal, Lingue Koto, Senegal Oriental | JN083622 | JN083753 | – | JN083512 | JN083682 |
| *Pterocarpus steinbachianus*  Harms | Klitgård 1465 (K) | Bolivia, Beni | KF436458 | KF436475 | KF436445 | KF436435 | – |
| *Pterocarpus ternatus*  Rizzini | Harley 26138 (K) | Brazil, Bahia | JN083629 | JN083761 | JN083575 | JN083521 | JN083689 |
| *Pterocarpus tinctorius*  Welw. | Delvaux 650 (K) | Democratic Republic of Congo, Katanga | JN083632 | JN083764 | JN083578 | JN083524 | JN083692 |
| *Pterocarpus tinctorius*  Welw. | Mwasumbi & Clarke 3608 (K) | Tanzania, Lindi District | JN083633 | JN083765 | JN083579 | JN083525 | JN083693 |
| *Pterocarpus tinctorius*  Welw. | Reekmans 9232 (K) | Burundi | JN083631 | JN083763 | JN083577 | JN083523 | JN083691 |
| *Pterocarpus tinctorius*  Welw. | Rees T151 (K) | Tanzania, T8, Yerende Ferry Selous Game Reserve | – | JN083767 | JN083581 | JN083528 | JN083696 |
| *Pterocarpus tinctorius*  Welw. | Schmitz 5970 (K) | Dem. Rep. Congo (Congo), Katanga, Terr. Jadotville | JN083630 | JN083762 | JN083576 | JN083522 | JN083690 |
| *Pterocarpus tinctorius*  Welw. | Wagemans 1769 (K) | Dem. Rep. Congo (Congo/Leopoldville), Boma, INEAC-Luki | JN083634 | JN083766 | JN083580 | JN083526 | JN083694 |
| *Pterocarpus villosus*  Mart. ex Benth. | Alencar 237 (K) | Brazil, Piauí | JN083636 | JN083769 | JN083582 | JN083530 | JN083698 |
| *Pterocarpus zehntneri*  Harms | Ribon 23137 (K) | Brazil, Espirito Santo | JN083637 | JN083770 | JN083583 | JN083531 | JN083699 |
| *Tipuana tipu* (Benth.)  Kuntze * | Klitgård 13 (AAU) | Brazil, Minas Gerais | AF189056 | JN083706 | AF270882 | - | - |

**Table S1.3:** Sampled vouchers, collector information and collection localities for *Pterocarpus* species used in the BEAST inference based on GenBank data. Herbaria from which accessions were collected are cited next to the collector name and number. GenBank names for each Sanger locus are given in the latter five columns. Loci and species for which GenBank data were not available are shown with a dash ‘-‘. Outgroup taxa from the Pterocarpus clade are marked with an asterisk (*).

| **Locus** | **JModeltest substitution model** | **BEAST substitution model** |
| --- | --- | --- |
| ITS | TrN + G | GTR + Gamma |
| *mat*K | TVM + G | HKY + Gamma + Invariant sites |
| *ndhF-rpl32* | TVM + G | HKY + Gamma + Invariant sites |
| *rbcl* | HKY + I + G | HKY + Gamma + Invariant sites |
| *trn*L*-trn*F | HKY + G | HKY + Gamma + Invariant sites |

**Table S1.4:** Substitution models used for inference of time-calibrated phylogenetic trees in BEAST 1.8.0. Column one shows the locus alignment, column two shows the model-testing output from Jmodeltest and column three shows the substitution model implemented in BEAST to best approximate the model selected by Jmodeltest.

**Table S1.5:** AIC tables and estimates for parameters for all six models tested in BioGeoBEARS for both realm and biome analyses. The best supported models, chosen using AICc, are emboldened.

*Realm*

| **Type** | **Model** | **LnL** | **Params** | **d** | **e** | **AICc** | **AIC wt** |
| --- | --- | --- | --- | --- | --- | --- | --- |
| **Trait independent** | *DEC* | -25.064 | 2 | 0.07 | 2.179x10^-8^ | 54.649 | 0.215 |
|  | ***DIVALIKE*** | **-24.131** | **2** | **0.078** | **7.207x10^-9^** | **52.785** | **0.545** |
|  | *BAYAREALIKE* | -34.536 | 2 | 0.086 | 0.248 | 73.594 | 1.651 x10^-5^ |
| Trait dependent | *Traits only*  *(1 switching rate)* | -84.646 | 1 | 1.00x10^-5^ | 1.00x10^-5^ | 171.459 | 2.34 x10^-26^ |
|  | *Traits only*  *(2 switching rates)* | -84.449 | 2 | 1.00x10^-5^ | 1.00x10^-5^ | 173.419 | 1.05x10^-26^ |
|  | *DEC_t12_t21_m2* | -34.904 | 5 | 0.041 | 1.00x10^-12^ | 82.808 | 1.72 x10^-6^ |

*Biome*

| **Type** | **Model** | **LnL** | **Params** | **a** | **AICc** | **AIC wt** |
| --- | --- | --- | --- | --- | --- | --- |
| **Trait independent** | ***‘Markov-k’*** | **-163.026** | **1** | **0.177** | **328.218** | **0.333** |
| Trait dependent | *‘Markov-k’_t12_t21_m2* | -174.192 | 3 | 0.177 | 355.475 | 4.02x10^-7^ |

**Figures**

**Figure S1.1:** **a)** Distribution map of all *Pterocarpus* species (blue points), along with WWF biogeographical realms (labelled zones) and terrestrial biomes (colours, shown in the legend), as defined in Olson et al. (2001), used for biogeographical analyses.

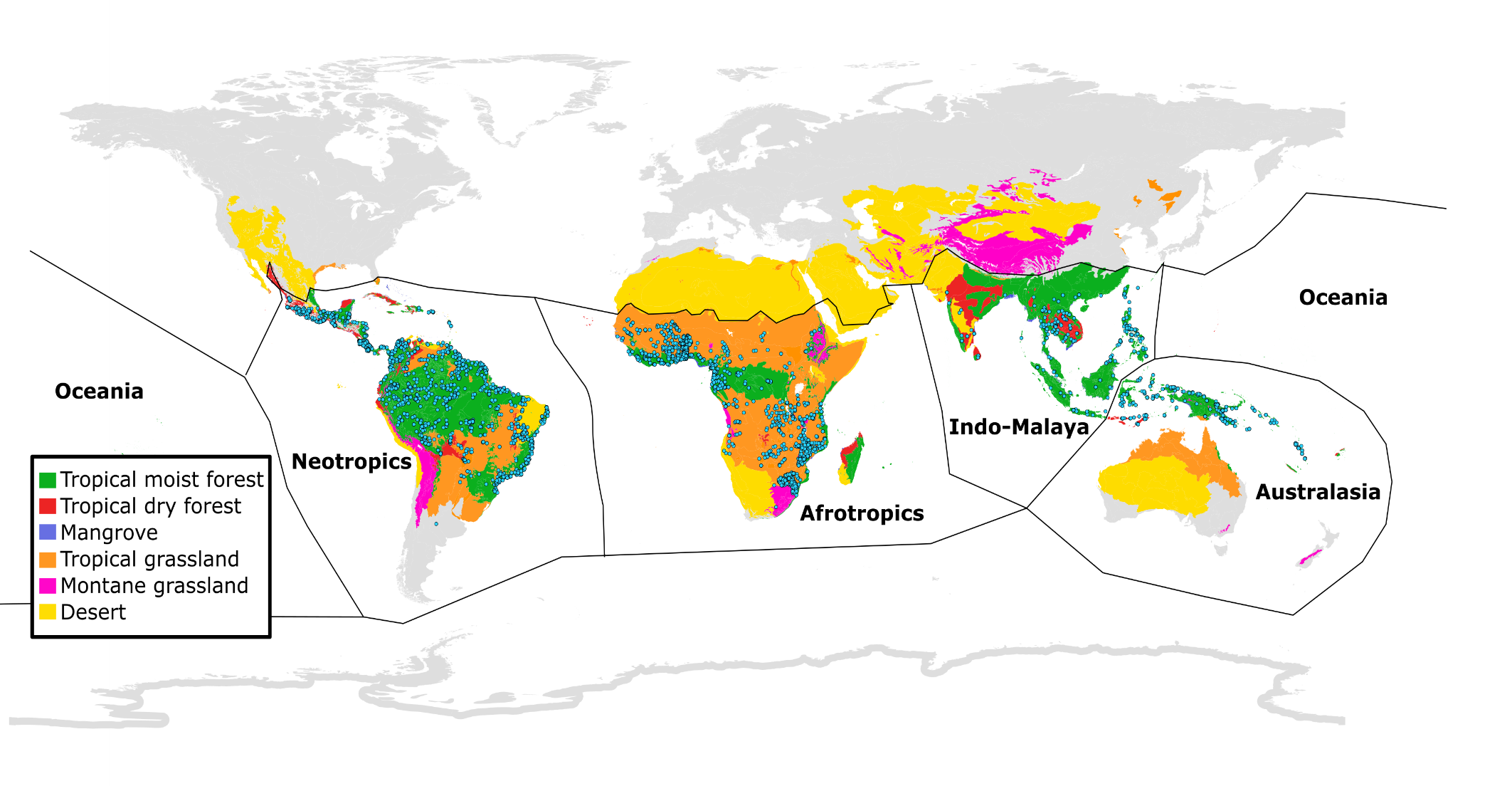

**b)** Fruit dispersal traits plotted on the ASTRAL tree along with the biogeographical realm and biome occupied by each species, based on classifications by Olson *et al.* (2001).

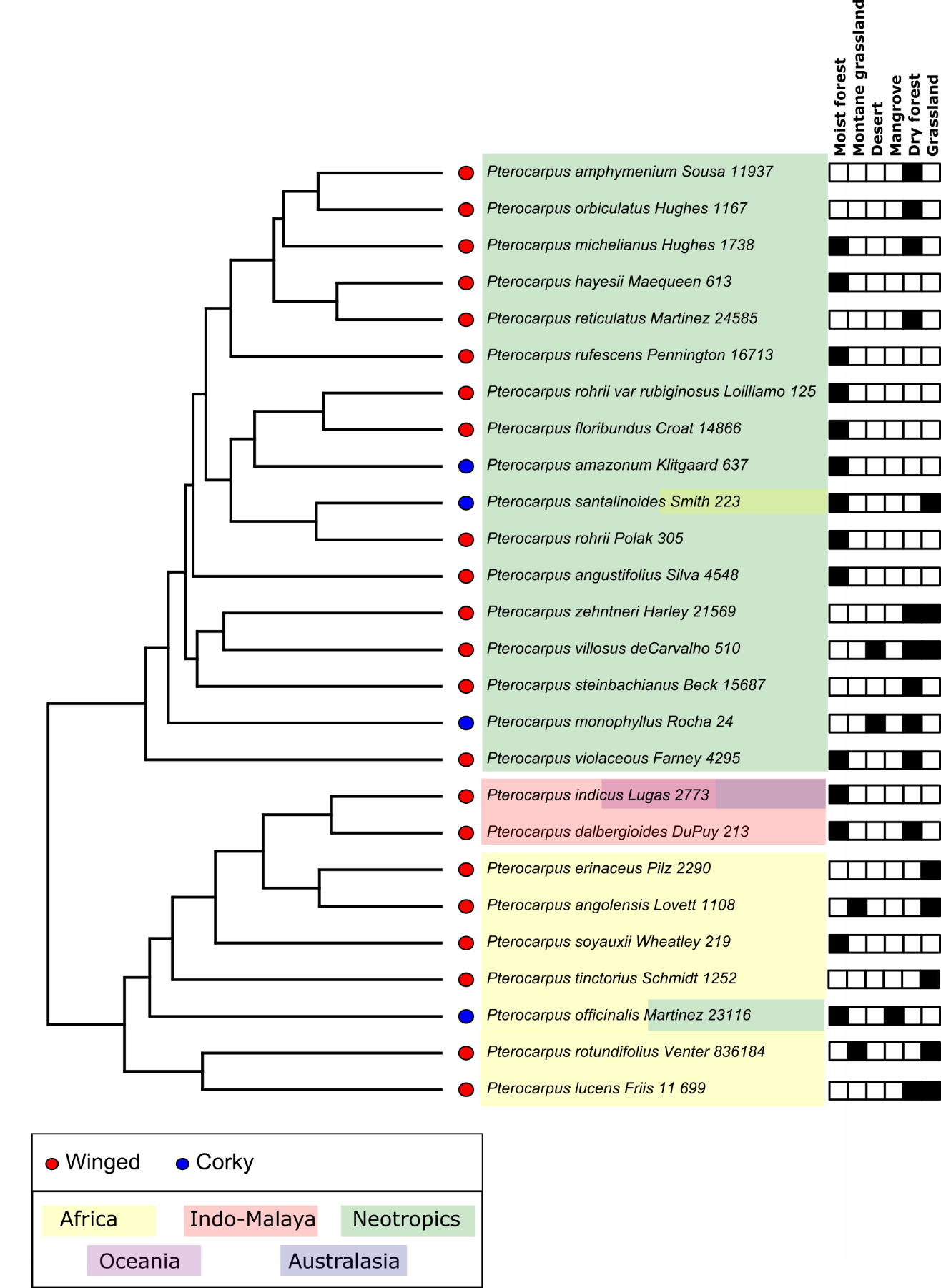

**Figure S1.2:** Gene recovery heatmap, indicating percentage of the reference protein length recovered for each locus in the Angiosperms-353 bait kit for each species in the phylogenomic dataset.

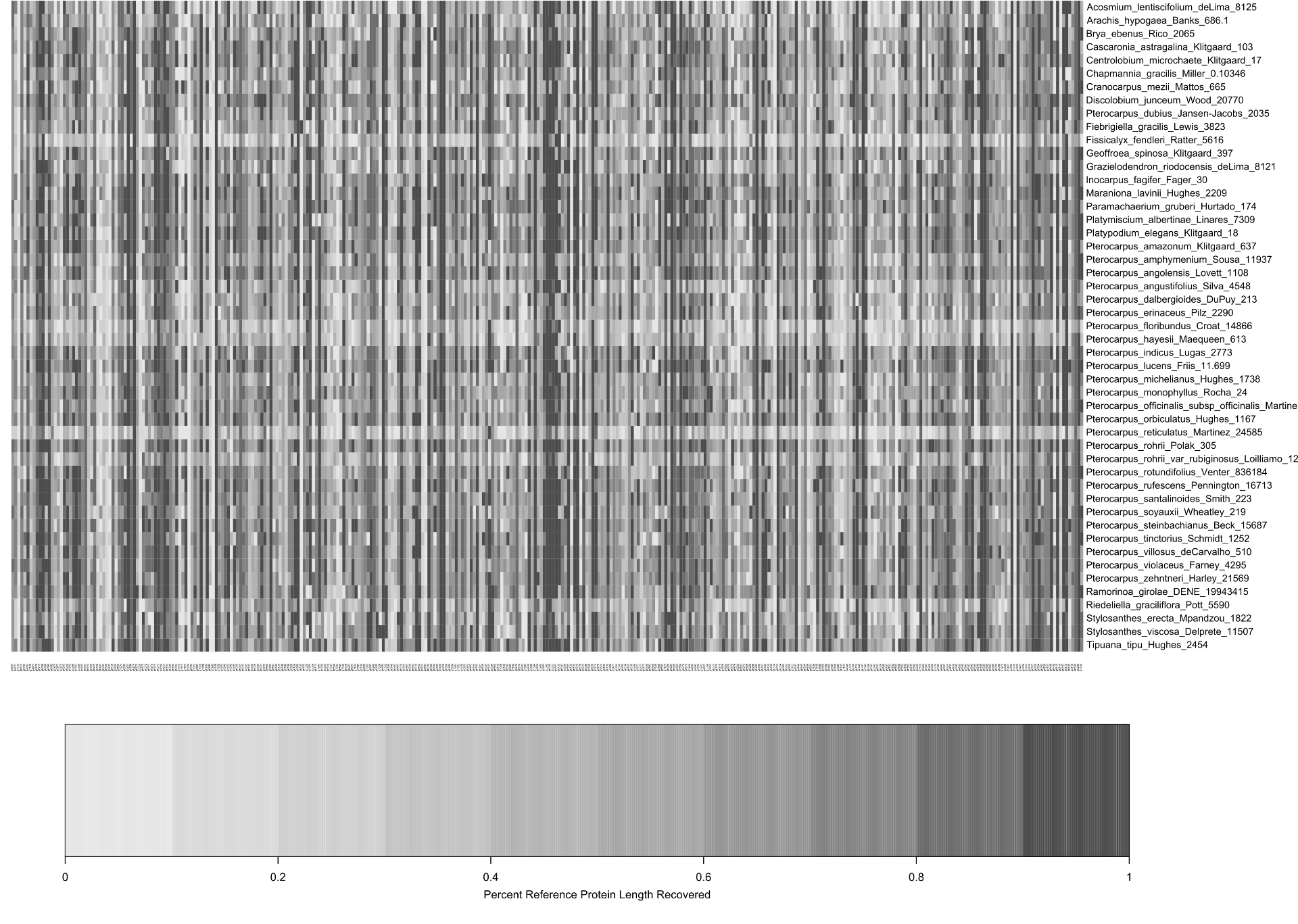

**Figure S1.3:** Phylogenetic tree of the Pterocarpus clade based on RAxML analysis of the concatenated, 303-locus phylogenomic alignment. Bootstrap values (BS) are shown for each node. The red and green shading indicates species belonging to the palaeotropical and neotropical subclades of *Pterocarpus*, respectively, and the extent of the genus *sensu stricto* is shown by the black bar. The dashed box around Pterocarpus dubius highlights its placement outside Pterocarpus s.str., revealing the paraphyletic nature of the genus. Photographs show winged fruits of *Pterocarpus erinaceus* (© Gwilym Lewis), coriaceous fruits of *Pterocarpus monophyllus* (*©* Domingos Cardoso) and winged fruits of *Pterocarpus violaceus* (© Reinaldo Aguilar, CC BY-NC-SA 2.0).

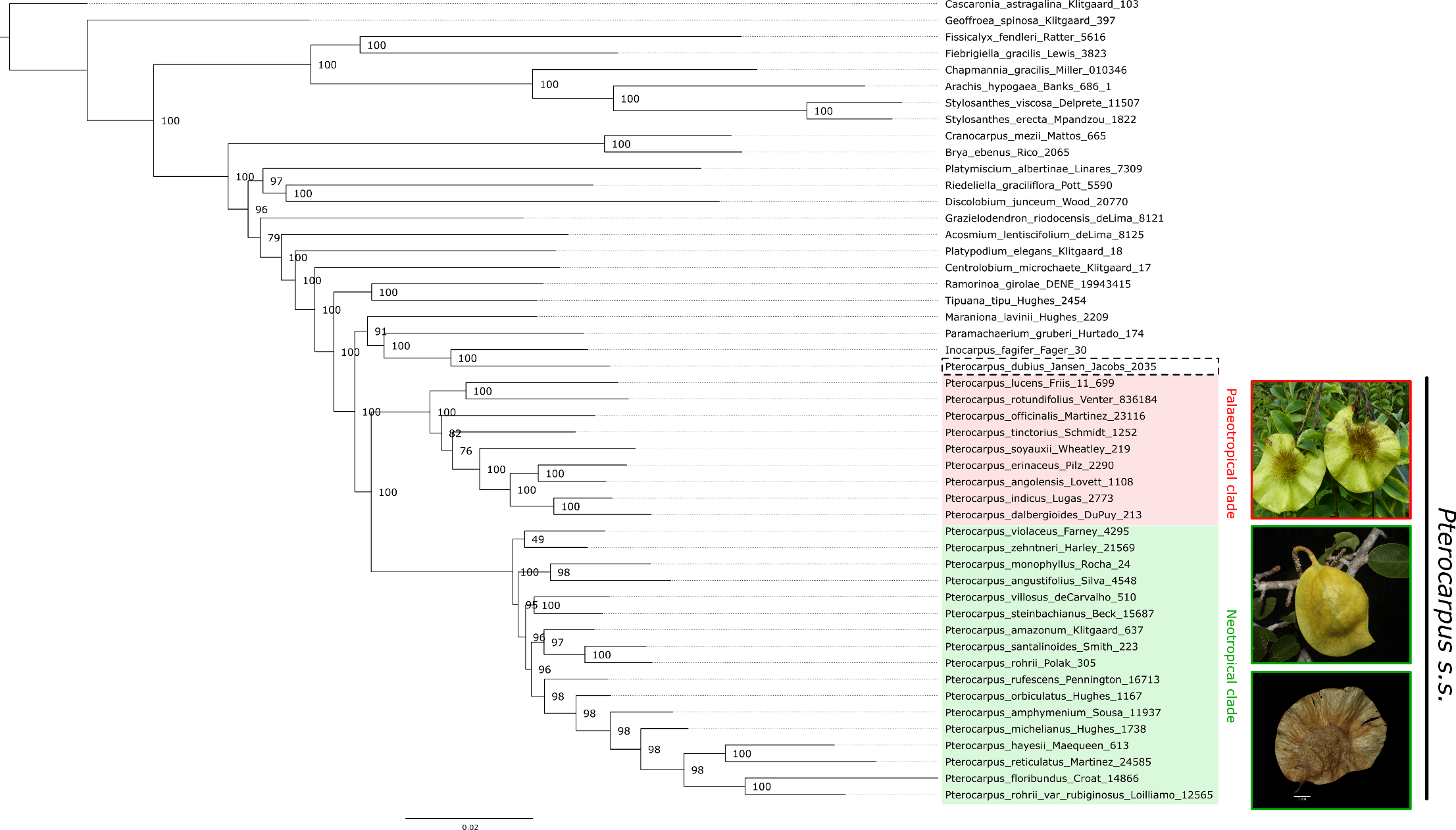

**Figure S1.4:**
**a)** Scatterplot of two MCMCtree runs, showing posterior date estimates for each node from the two runs. Each point lies on the line X=Y, indicating perfect correlation between posterior estimates and so MCMC convergence between runs.

**b)** MCMC traces of both runs showing likelihood at each generation

**c)** Distribution of posterior and prior estimates from one run of MCMCtree

**a)**

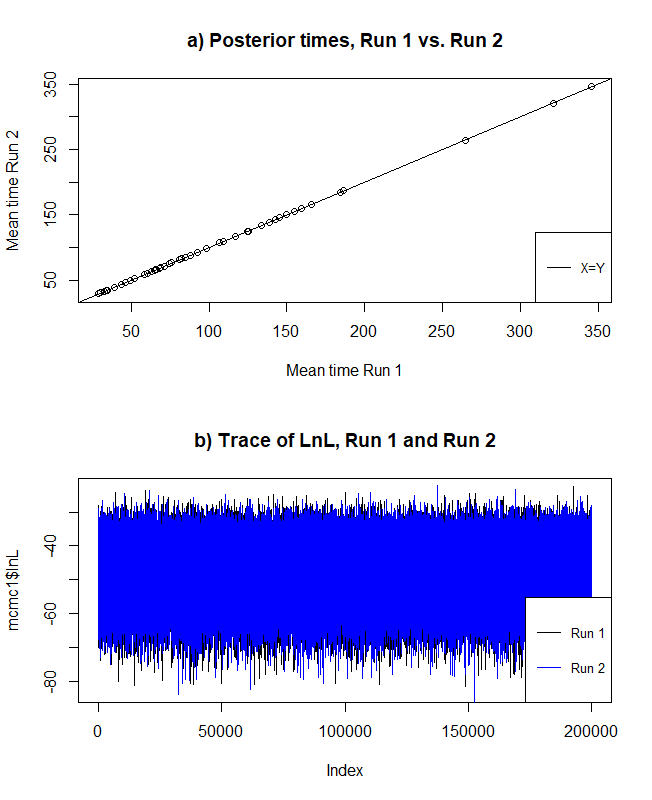

**b)**

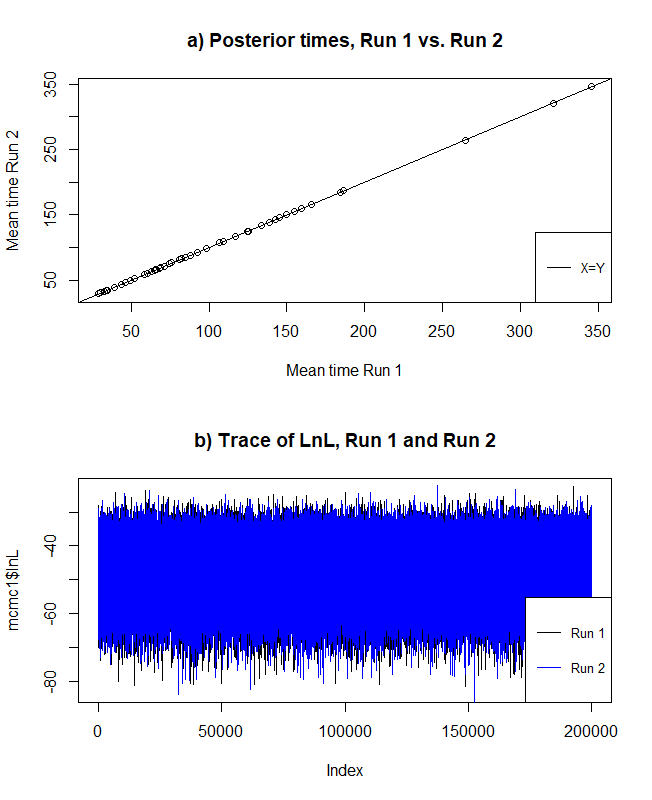

**c)**

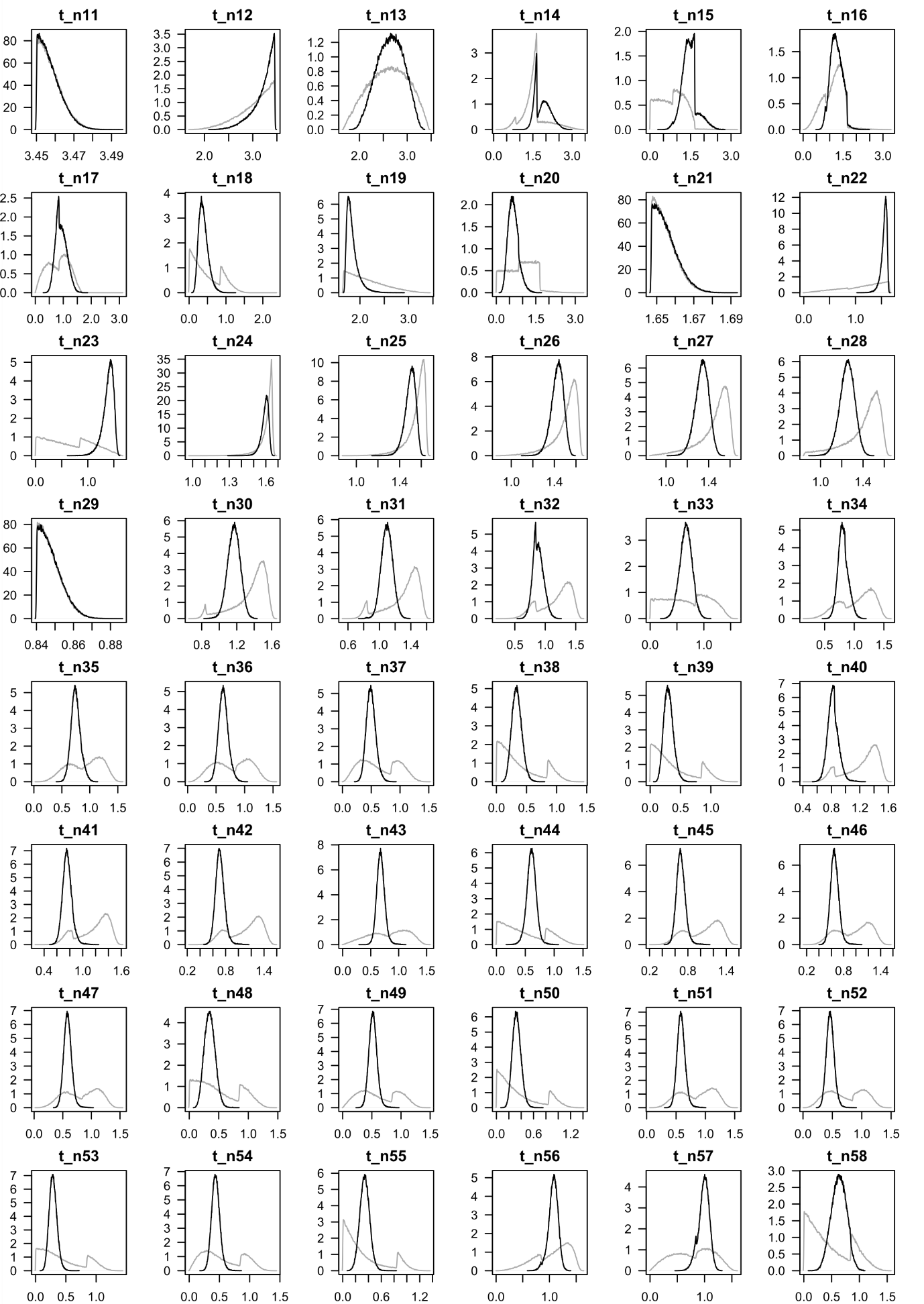

**Figure S1.5:** Densitree, showing distribution of output trees from MCMCtree based on the phylogenomic dataset. Shading on the background of the phylogram represents geological epochs, which are labelled beneath the tree. The red and green shading indicates species belonging to the palaeotropical and neotropical subclades of *Pterocarpus*, respectively, and the extent of the genus *sensu stricto* is shown by the black bar. The dashed box around *Pterocarpus dubius* highlights its paraphyly with the rest of the genus.

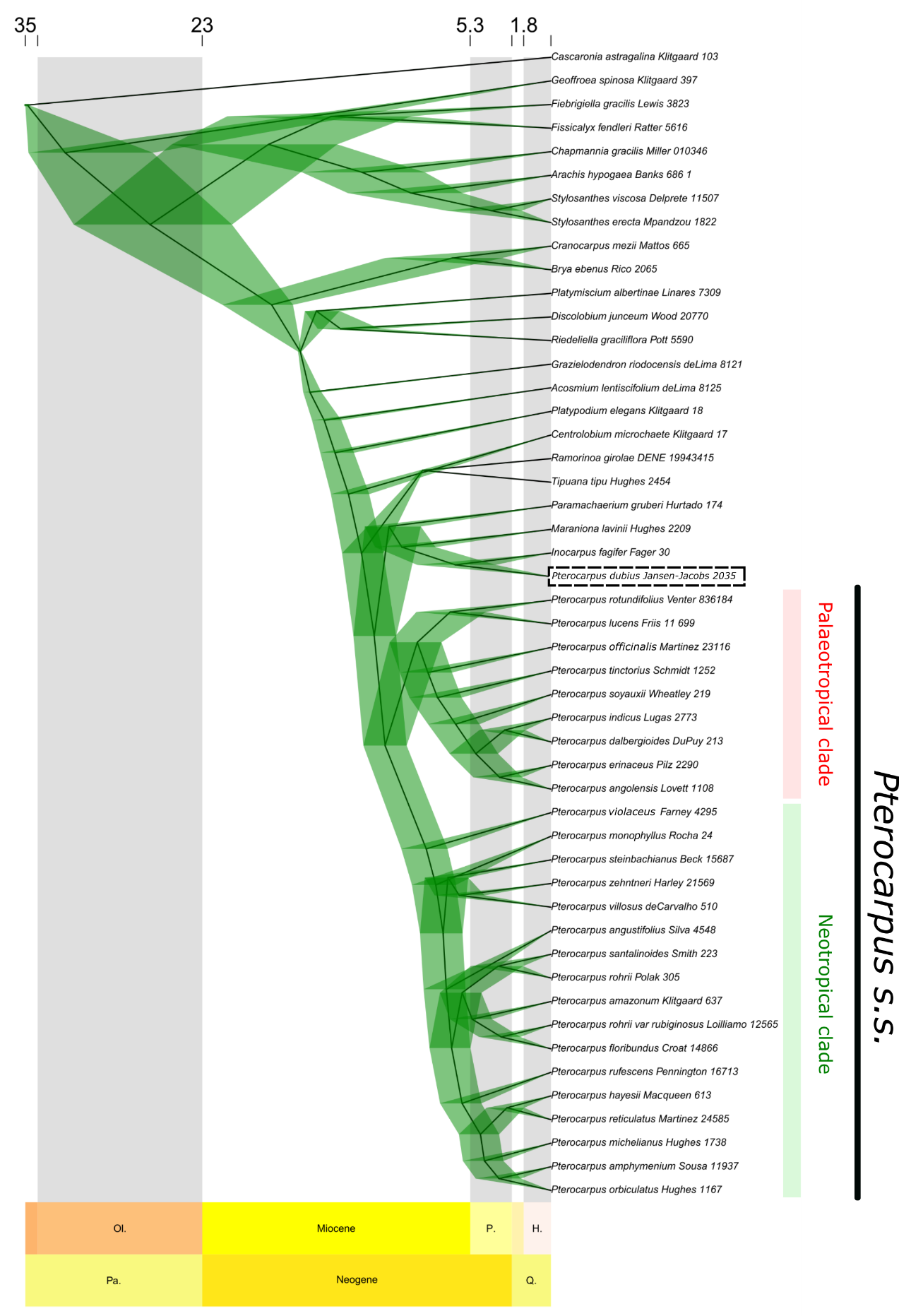

**Figure S1.6:** BEAST tree inferred from the one nuclear and four chloroplast loci downloaded from GenBank for each *Pterocarpus* species and outgroups.

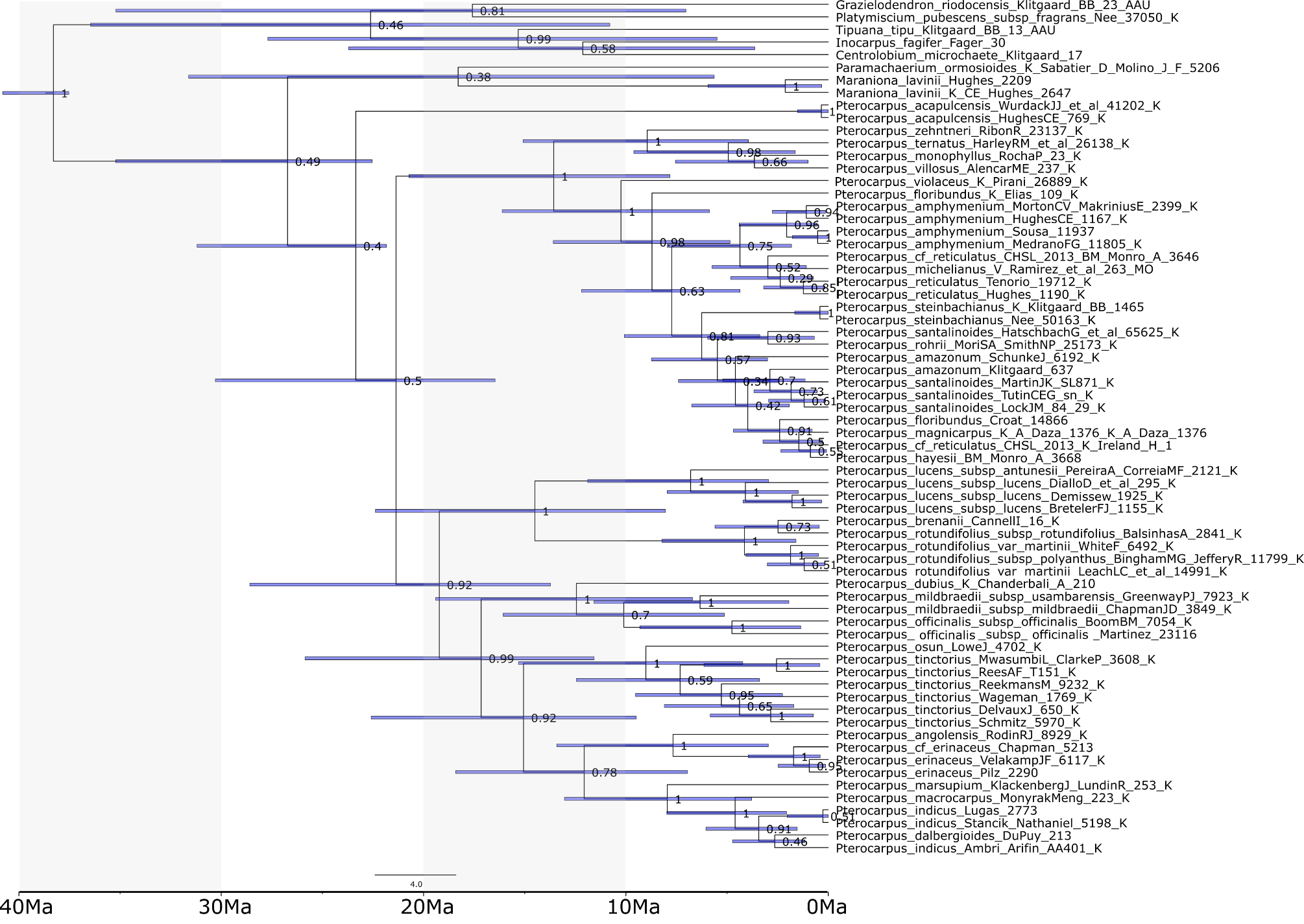

**Supplementary Methods.**

*BEAST analysis*

For comparison to the time-calibrated phylogenomic tree inferred with MCMCtree, Sanger sequencing data downloaded from GenBank were used to perform divergence time estimation in BEAST v.1.8.0 (Drummond & Bouckaert, 2015) on the CIPRES web portal. Multiple runs of BEAST were undertaken using several different tree models (Yule, birth-death and birth-death (incomplete sampling)) to compare model fit. Substitution rate models were chosen for each of the five loci using AICc comparisons implemented in JModelTest (Darriba et al., 2012). Loci and their corresponding substitution rate models are outlined in Appendix S1, Table S1.4. Tree model choice was based on path sampling analysis output from BEAST, using 10 million generations with 30 path steps, resulting in the use of the Birth-Death model for inference. Dated trees were generated using a relaxed lognormal clock model, informed by coefficient of variation values >0.1, taken from the BEAST log file. Two of the three fossil calibration points in the MCMCtree phylogenomic dating analysis were used to provide date estimates in BEAST (Table1).

Phylogenetic inference in BEAST comprised three independent runs of 50 million generations (giving a total of 150 million generations), which were combined using Logcombiner v.1.8.2 with a burn-in level of 10%. Convergence of runs was assessed using effective sample size (ESS) of all parameters in Tracer v. 1.6 with a threshold value of 200. The final ultrametric trees were generated in TreeAnnotator v1.8.2 and visualised using FigTree v1.4.4 (Rambaut, 2014).

*GBIF data collection and cleaning*

Geographical distribution data were collected from GBIF ([www.GBIF.org](http://www.gbif.org) (06 November 2019), GBIF Occurrence Download <https://doi.org/10.15468/dl.tr3h2y>) based on the curated taxonomic name list described above. Only records corresponding to ‘preserved specimens’ and ‘material samples’ were included in this dataset, and all ‘cultivated’ specimens were removed. Moreover, cultivars of the peanut (*Arachis hypogaea*), which is part of the Pterocarpus clade, were removed to prevent the inclusion of any cultivated specimens in our geographical dataset. This dataset was then further refined using the R package ‘CoordinateCleaner’ ((Zizka et al., 2019); <https://github.com/ropensci/CoordinateCleaner>) and the input for our *BioGeoBears* analysis was produced from these data with Alex Zizka’s ‘biogeography in R’ scripts (<https://github.com/azizka/Using_biodiversity_data_for_biogeography>). These records and the specimens underpinning them were then examined in detail by taxonomic experts on the *Pterocarpus* clade (B. B. Klitgaard and G. P. Lewis) to further remove records which were mis-identified.

*Assigning species to realms and biomes*

From these refined data, species were assigned to biogeographical regions using the *wwfLoad()* function in ‘speciesgeocodeR’. Coordinates for each species were plotted on two maps: the first was based on the WWF terrestrial realms, representing five major biogeographical realms within the tropics (Australasia, Afrotropics, IndoMalaya, Neotropics and Oceania (Olson et al., 2001)). The second was based on the WWF terrestrial biomes map, representing six of the world’s tropical terrestrial biomes (tropical moist forest (i.e., ‘rainforest’), montane grassland, desert, mangrove, tropical dry forest and tropical grassland (i.e., ‘savanna’) (Olson et al., 2001)). These regions are displayed in Appendix S1, Fig. S1.1a. This allowed us to extract both the biogeographical area and the biome in which each sample was found, which then allowed us to assign each species to one or more biogeographical regions and biomes. Finally, biogeographical assignments for each *Pterocarpus* species were cross-referenced with monographs and previous studies of *Pterocarpus* (Rojo, 1972, Saslis-Lagoudakis et al., 2011, Klitgård et al., in prep), and with the Plants Of the World Online database (<http://powo.science.kew.org/>). All traits (fruit morphology, realm assignment and biome assignment) are visualised onto the *Pterocarpus* phylogenetic tree in Appendix S1, Fig. S1.1b.

*Biogeographical analysis*

Trait-dependent historical biogeography analyses were performed using *‘*BioGeoBEARS*’* to assess the effect of dispersal traits on the biogeographical history of *Pterocarpus.* The fit of trait-dependent biogeographical models outlined by Klaus and Matzke (2020) and non-trait-dependent models were assessed for both biogeographical realms and biomes independently. For realms, six models were tested, all of which incorporated dispersal, extinction and range switching. First, we tested between the standard models implemented in ‘BioGeoBEARS’, which are DEC (Ree & Smith, 2008), which additionally accounts for vicariance and ‘subset’ sympatric speciation, DIVALIKE (Ronquist, 1997), which also accounts for vicariance, and BAYAREALIKE (Landis et al., 2013), which additionally accounts for widespread sympatric speciation but not vicariance. For biomes, we assessed the fit of Markov-*k* (Mk) models (Lewis, P. O., 2001) (i.e., BAYAREALIKE a+, d=e=0 in BioGeoBEARS) to accommodate a standard unordered character with equal rates of character evolution, after (Kriebel et al., 2019). Markov-*k* more accurately models the inheritance of ecological preferences in daughter species and avoids biases towards reconstructing large ancestral ranges when compared to other biogeographical models such as DEC.

In addition to these standard models, we also included four trait-dependent models in our comparisons. The first of these models parameterised only traits with one rate of switching (‘*Trait_1Rate’*, which describes switching from trait 1 to trait 2 (t_12_)). The second was ‘*Trait_2Rates’*, which describes two rates of trait switching (t_12_ and t_21_). For realm, the third was *DEC_t12_t21_m2*, which is a trait-dependent extension of the DEC model that describes two rates of trait switching as well as a multiplier on the base anagenetic dispersal rate *d* (known as ‘*m2’*). Finally, for biome the fourth model was a trait-dependent implementation of the Markov-*k* model (‘*Markov-k’_t12_t21_m2*).

The model which was best described by the data was chosen using AICc and Akaike weights, from which ancestral ranges and range shifts were estimated across *Pterocarpus*. Ancestral ranges were plotted onto the time-calibrated phylogenomic tree inferred with MCMCtree using the function ‘*plot_BioGeoBears_results()*’ for realm and biome independently. For realm, the maximum number of range areas was set to five, and for biome it was set to six, as these were the total number of biogeographical areas used for each analysis. Fixed parameters in each model were given ‘initial’ and ‘estimate’ values from ‘0.0’ to ‘0.00001’ as per the recommendations of the ‘BioGeoBEARS’ authors. To examine patterns of biome shifting across the *Pterocarpus* tree, we then counted the number of transitions between the most likely ancestral states at each node of Fig. 2b, and produced a matrix of these shifts. This matrix was plotted using the *plotmat* function in the R package *diagram* (Soetaert, 2012).
